# Alzheimer mutations stabilize synaptotoxic γ-secretase-substrate complexes

**DOI:** 10.1101/2023.09.08.556905

**Authors:** Sujan Devkota, Rui Zhou, Vaishnavi Nagarajan, Masato Maesako, Hung Do, Arshad Noorani, Caitlin Overmeyer, Sanjay Bhattarai, Justin T. Douglas, Anita Saraf, Yinglong Miao, Brian D. Ackley, Yigong Shi, Michael S. Wolfe

## Abstract

Alzheimer’s disease is characterized pathologically by cerebral deposition of 42-residue amyloid β-peptide (Aβ42), proteolytically produced from amyloid precursor protein (APP) by β- and γ-secretases.^1^ Although mutations in APP and presenilin, the catalytic component of γ-secretase, cause familial Alzheimer’s disease (FAD), a role for Aβ42 as the primary disease driver has not been clearly established and remains controversial.^2,3^ Here we show through comprehensive analysis of the multi-step proteolysis of APP substrate C99 by γ-secretase that FAD mutations are consistently deficient in early proteolytic events, not later events that produce secreted Aβ peptides. Cryo-electron microscopy revealed that a substrate mimetic traps γ-secretase at the transition state for intramembrane proteolysis, and this structure closely aligns with activated enzyme-substrate complex captured by molecular dynamics simulations. *In silico* simulations and fluorescence lifetime imaging microscopy in cultured cells support stabilization by FAD mutations of enzyme-substrate and/or enzyme-intermediate complexes. Neuronal expression of C99 and/or presenilin-1 in *Caenorabditis elegans* led to age-dependent synaptic loss only when one of the transgenes carried an FAD mutation. Designed mutations that stabilize the enzyme-substrate complex and block proteolysis likewise led to synaptic loss. Collectively, these findings implicate the stalled process—not the released products—of γ-secretase cleavage of substrates in FAD pathogenesis.

---

The discovery of dominant missense mutations in APP associated with FAD led to original formulation in 1991 of the amyloid hypothesis,^4,5^ which posits that aggregation of secreted Aβ peptides, particularly Aβ42, leads to a cascade of events culminating in neurodegeneration and dementia. Subsequent findings that presenilins are sites of FAD mutations that alter Aβ production, are essential for γ-secretase processing of APP to Aβ, and comprise the catalytic component of the γ-secretase complex provided strong support for the amyloid hypothesis.^1^ Nevertheless, the assembly state of neurotoxic Aβ42 and associated signaling pathways remain unclear,^6^ and clinical candidates targeting Aβ or its aggregates have shown little or no benefit in the prevention or treatment of Alzheimer’s disease,^7,8^ the recent approval of anti-Aβ monoclonal antibodies notwithstanding,^8,9^ raising doubts about Aβ42 as a major driver of the disease process.

The pathology, presentation and progression of FAD are closely similar to those of the more common sporadic late-onset Alzheimer’s disease,^10,11^ and the dominantly inherited monogenic nature of FAD suggests that elucidation of pathogenic mechanisms should be more tractable. Because dominant missense FAD mutations are only found in the substrate and the enzyme that produce Aβ, such mutations all likely lead to altered proteolytic processing of APP substrate by γ-secretase. However, this processing is complex, with the APP transmembrane domain (TMD) cleaved multiple times by the membrane-embedded γ-secretase complex to produce Aβ peptides along two pathways: Aβ49→Aβ46→Aβ43→Aβ40 and Aβ48→Aβ45→Aβ42→Aβ38 (Fig. 1a).^12^ We recently reported comprehensive analysis of effects on each of these proteolytic events for 14 FAD mutations in the APP TMD, finding that every mutation was deficient in the first or second carboxypeptidase trimming step, elevating levels of Aβ peptides of 45 residues and longer.^13^ Such complete and quantitative analysis has not been reported for any presenilin FAD mutations.

**Fig. 1.**
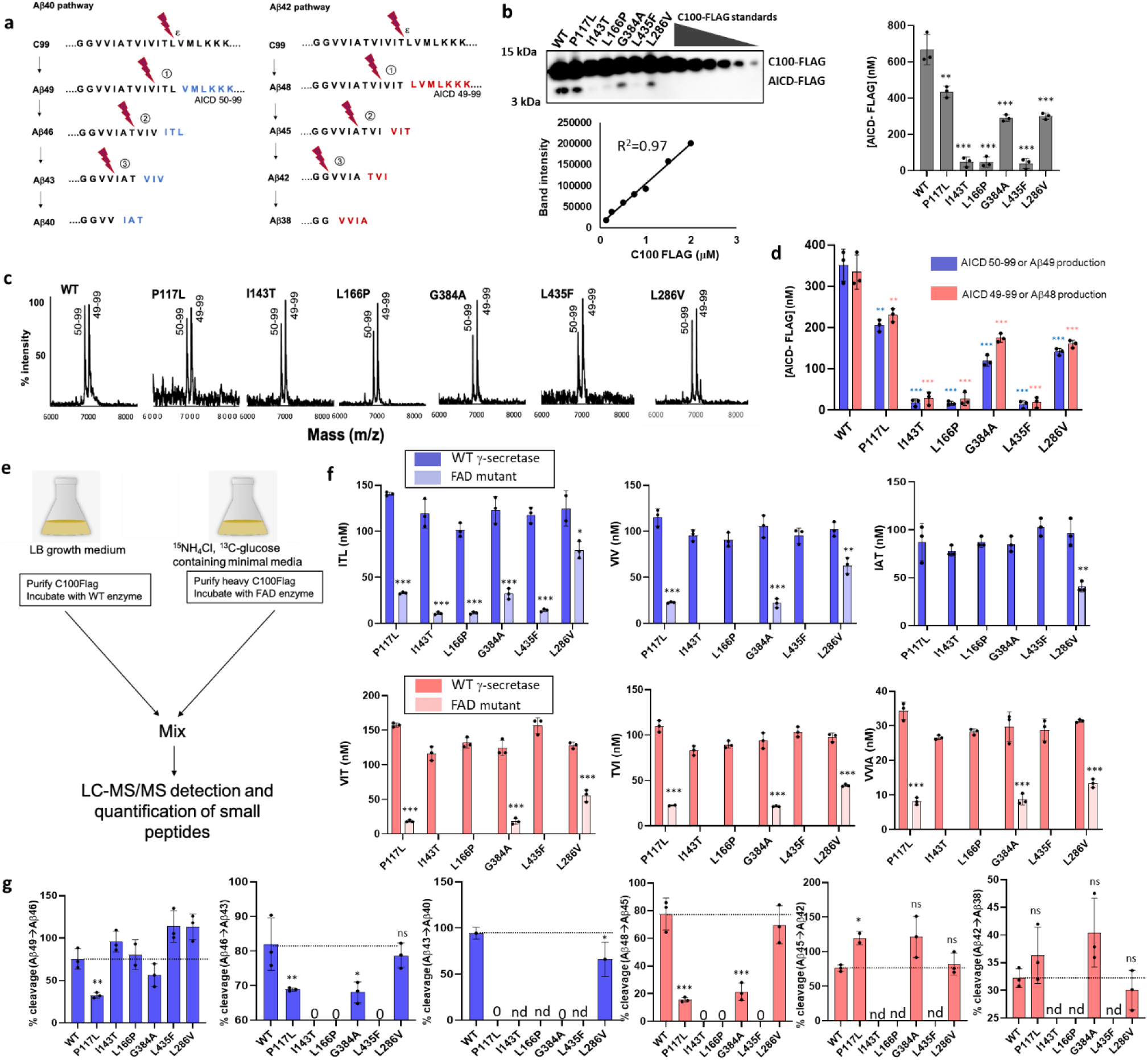
Analysis of all co-products generated from processive proteolysis of APP substrate by γ-secretase reveal reduction of early proteolytic events with six FAD PSEN1 mutations. **a**, Schematic of proteolytic processing of APP substrate by γ-secretase. **b**, Quantitative western blotting of total AICD, using purified substrate C100Flag to generate a standard curve for densitometry. **c**, MALDI-TOF analysis shows ratios of peak heights of AICD49-99/AICD50-99 produced by γ-secretase with WT versus six FAD mutants of PSEN1. **d**, Quantification of AICD49-99 and AICD50-99 production, corresponding to Aβ48 and Aβ49 production, respectively. **e**, Schematic of light-vs. heavy-isotope labeling of APP substrate for LC-MS/MS analysis of effects of PSEN1 FAD mutations on carboxypeptidase trimming steps. **f**, Bar graphs of small peptide coproduct for each trimming step. Blue graphs: first, second and third trimming step for the Aβ49→Aβ40 pathway. Red graphs: trimming steps for the Aβ48→Aβ38 pathway. Dark and light bars: coproduct formation from WT and FAD-mutant γ-secretase, respectively. **g**, Bar graph of percent cleavage efficiencies of each trimming step relative to WT enzyme. Where the level of precursor Aβ peptide for a given trimming step is zero, the efficiency of this cleavage event could not be determined (nd). In all graphs, n=3, unpaired two-tailed T-test compared FAD mutant to WT, *p≤ 0.05, **p≤ 0.01, ***p≤ 0.001.)

Six FAD mutations in presenilin-1 (PSEN1) were separately installed into a tetracistronic pMLINK construct that encodes all four components of the γ-secretase complex,^14^ in a multi-step process starting from monocistronic pMLINK encoding human PSEN1 (Extended data, Fig. 1). Wild-type (WT) and FAD-mutant γ-secretase complexes were expressed in and purified from suspension human embryonic kidney (HEK) 293 cells as previously described.^13^ Each purified protease variant (30 nM) was incubated with saturating levels (3 µM) of APP-based recombinant substrate C100-Flag at 37 °C for 16 h, when the rate of substrate cleavage was still in the linear range. Rates of formation of APP intracellular domain (AICD) coproducts AICD50-99 and AICD49-99 (Fig. 1a)—produced through initial endoproteolytic (ε) cleavage—were quantified in two steps: quantitative western blotting of total Flag-tagged AICD product, using C100-Flag to establish a standard curve (Fig. 1b), and matrix-assisted laser desorption/ionization-time of flight mass spectrometry (MALDI-TOF MS) to determine the ratio of the two specific AICD products (Fig. 1c). All six FAD-mutant enzymes skewed ε cleavage (Fig. 1c) toward AICD49-99 and therefore toward the Aβ48→Aβ42 pathway, as previously reported for PSEN1 FAD mutations.^15^ All six mutant proteases were also deficient in ε cleavage, with lower rates of production of total AICD, AICD50-99 and AICD49-99 than was seen with WT γ-secretase (Fig. 1b,d), which is consistent with previous reports of reduced ε proteolysis with PSEN1 FAD mutations.^16–18^ FAD-mutants P117L, G384A and L286V produced ∼30-50% less AICD, while I143T, L166P and L435F were >90% deficient in ε proteolysis. Quantification of AICD49-99 and AICD50-99 (Fig. 1d) provided indirect quantification of the production of respective ε cleavage coproducts Aβ48 and Aβ49, data that was used below to determine efficiencies of subsequent carboxypeptidase trimming steps.

To quantify the production of each small peptide coproduct of processive carboxypeptidase cleavage by WT and FAD-mutant γ-secretase complexes, APP-based recombinant substrate C100-Flag was expressed in and purified from *E. coli* under conditions that would provide either “light” (^12^C/^14^N) or “heavy” (^13^C/^15^N) isotopic labeling (Extended data, Fig. 2). “Light” substrate was incubated with WT γ-secretase, while in parallel “heavy” substrate was incubated with FAD-mutant proteases (Fig. 1e). This allowed subsequent mixing of equal volumes of WT and FAD-mutant enzyme reaction mixtures, with detection and direct comparison of tri- and tetra-peptide proteolytic coproducts from the respective reactions through liquid chromatography coupled with tandem mass spectrometry (LC-MS/MS).^12,13^ The three most abundant fragment ions of these small peptides were quantified, first using purified synthetic peptides to establish standard curves and then for the mixed 1:1 enzyme reactions with “light” and “heavy” substrates. Separately incubating light and heavy C100Flag substrate with WT γ-secretase, followed by mixing the two samples before LC-MS/MS analysis validated the method, giving equal levels of light and heavy tri- and tetrapeptide coproducts (Extended data, Fig. 3). WT γ-secretase also produced consistent levels of each peptide in the LC-MS/MS runs when mixed with the different FAD-mutant enzyme reactions (Fig. 1f), providing internal standardization to measure the effect of FAD mutation.

Due to the reduced ε proteolytic activity seen with all six mutant protease complexes, formation of each small peptide coproduct was also lower than that produced from the WT enzyme (Fig. 1f). Together with the quantification of AICD coproducts, quantification of these small peptides provided the degree of degradation and production of each Aβ product and therefore the percent efficiency of each carboxypeptidase trimming step for WT versus FAD-mutant enzyme (Fig. 1g). Five of the six FAD-mutant γ-secretase complexes (with L286V the exception) were deficient in Aβ48→Aβ45 trimming. Reduced Aβ49→Aβ46 trimming was seen with the P117L mutation, while Aβ46→Aβ43 trimming was strongly reduced with I143T, L166P and L435F mutations, and Aβ43→Aβ40 was reduced with P117L, G384A and L286V mutations. The degree of degradation and production for each Aβ peptide also allowed determination of the levels of each Aβ peptide from the enzyme reactions (Extended data, Fig. 4). As an additional cross-check of the LC-MS/MS method, the level of detected total AICD was found to be virtually identical with the sum of all calculated Aβ peptide levels produced from each enzyme reaction. Moreover, levels of AICD49-99 and AICD50-99 equaled the sum of Aβ peptides produced from their respective pathways (Extended data, Fig. 4). We have reported similar equimolar production of Aβ and AICD from the processing of 14 FAD-mutant APP substrates by WT γ-secretase.^13^ These results show that all six FAD PSEN1 mutations led to reduction in early proteolytic processing steps on APP substrate by γ-secretase (ε cleavage, Aβ49→Aβ46, and/or Aβ48→Aβ45 trimming), similar to results with 14 FAD mutations in APP substrate. Specific ELISAs also showed that while all six PSEN1 FAD mutations produce much less Aβ40 and Aβ42 compared to that produced from WT γ-secretase, Aβ40 is consistently reduced more than is Aβ42, thereby increasing the Aβ42/Aβ40 ratio (Extended Data, Fig. 5).

Because FAD mutations alter proteolytic processing of the APP TMD by γ-secretase, we sought a structural dynamic understanding of how the enzyme carries out intramembrane proteolysis. Toward this end, we designed and synthesized a series of full TMD substrate-based peptidomimetics as probes to trap the γ-secretase complex at the transition state for structure elucidation by cryo-electron microscopy (cryo-EM).^19,20^ These TMD substrate mimetics were comprised of a hydroxyethylurea-based transition-state analog,^21^ which targets the aspartyl protease active site of γ-secretase on PSEN1, linked to an APP TMD-based peptide containing helix-inducing α-aminoisobutyric acid (Aib) residues,^22^ which targets a proximal substrate-binding exosite (Fig. 2a).^23^ Both peptidomimetic components on their own are moderately potent inhibitors of γ-secretase (50% inhibitory concentrations, or IC_50_s, in the range of 40-60 nM toward 1 nM enzyme).

**Fig. 2.**
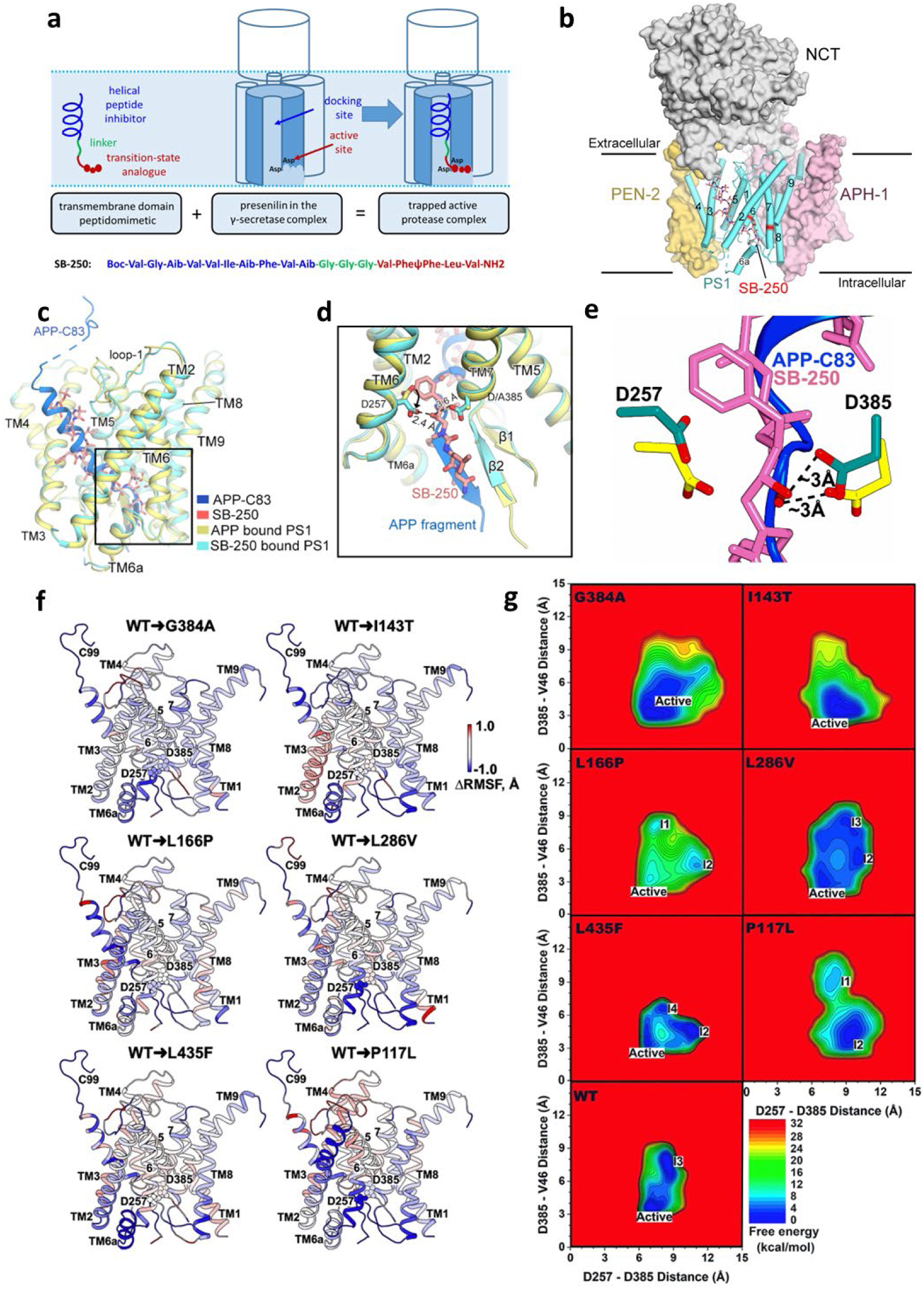
Structural analysis of active γ-secretase bound to a potent transmembrane substrate mimetic inhibitor vis-à-vis molecular dynamics simulation of activated substrate-enzyme complex. **a**, Schematic of substrate-based structural probe design and specific probe SB-250. **b**, The overall cryo-EM structure of SB-250 bound to WT γ-secretase complex. SB-250 is shown in sticks. **c**, The conformation of active PS1 (cyan) in complex with SB-250 (red) shows high similarity with catalytically inactive PS1 (yellow) covalently crosslinked to APP-C83 (marine). Superimposition of these two PS1 molecules reveals an r.m.s.d. (root-mean-square distance) of 0.428 Å over 250 Cα atoms. **d**, The structural probe traps active γ-secretase at the transition-state of intramembrane proteolysis. Note the rotation about the Cα-Cβ bond of catalytic residue D257 in SB-250-bound active γ-secretase compared to APP-C83-bound inactive enzyme (see curved arrow), positioning the aspartyl carboxylate for coordination with the transition-state-mimicking hydroxyl group of SB-250. **e**, Overlap of the cryo-EM structure of substrate mimetic probe bound to γ-secretase with the activated E-S complex from GaMD simulations reveals close similarity of the alignment and orientation of the active site aspartates, with respect to each other as well as with substrate/mimetic. Catalytic D257 and D385 are green for cryoEM structure and yellow for GaMD simulation. **f**, GaMD simulations of γ-secretase bound to C99 substrate show overall reduced flexibility with FAD PSEN1 mutations, implying complex stabilization. Root-mean-square fluctuations (RMSFs) of PSEN1 and C99 substrate within E-S complexes were calculated by averaging the RMSFs from individual MD simulations of each γ-secretase system. Changes in RMSFs (ΔRMSF) from WT to FAD-mutant PSEN1 were calculated by subtracting RMSFs of the WT E-S complexes from those with FAD-mutant PSEN1. **G**, Free energy profiles of WT and six FAD PSEN1 mutations from GaMD simulations of the Aβ49→Aβ46 trimming step.

However, linking them through short 9- or 10-atom spacers resulted in inhibitory potencies approaching stoichiometric levels (IC_50_ of 0.5 nM with 1 nM enzyme).^19,20^ Several of these TMD mimetic inhibitors were selected for study with γ-secretase by cryo-EM, and one of these compounds (**SB-250** in Fig. 2a), with a triglycine linker, provided an atomic-resolution (2.6 Å) structure (Fig. 2b-d; Extended Data Fig. 6 and Extended Data Table 1).

The structure of TMD mimetic **SB-250** bound to γ-secretase (this study) is remarkably similar to that of the enzyme bound to APP substrate^24^ (Fig. 2c). In both structures, the N-terminal region of the substrate/mimetic TMD is in a helical conformation and enveloped by PSEN1 TMDs, while the C-terminal region is in an extended conformation that interacts with the active site. However, the protease structure with bound APP substrate was accessed by disulfide crosslinking this substrate to PSEN1 through cysteine mutations as well as using catalytically inactive protease complex, with one of the two PSEN1 TMD aspartates mutated to alanine.^24^ In contrast, mimetic **SB-250** is bound noncovalently to the active γ-secretase complex, allowing coordination of the transition state-mimicking moiety with the two catalytic aspartates (Fig. 2d). This coordination, within hydrogen-bonding distance, between the mimetic hydroxyl group and the two PSEN1 TMD aspartates, is closely similar to what has been observed in x-ray crystal structures of water-soluble aspartyl proteases bound to related transition-state analogues.^25–27^ Thus, the new structure of TMD substrate mimetic **SB-250** bound to γ-secretase likely reveals the conformation of the enzyme-substrate (E-S) complex at the transition state, poised as it would be for intramembrane proteolysis.

In parallel with the development and analysis of structural probe peptidomimetics, we also used an *in silico* molecular dynamics system to understand γ-secretase action and the effects of FAD mutations. We recently reported the development of such models, for ε cleavage and the Aβ49→Aβ46 trimming step using all-atom Gaussian-accelerated molecular dynamics (GaMD) simulations.^28–30^ In these simulations, water entered the active site and coordinated between the two aspartates, with positioning for nucleophilic attack of the water oxygen on the carbonyl carbon of the scissile amide bond Leu49-Val50. Meanwhile, the carboxylic acid side chain of the protonated aspartate interacted with the oxygen atom of the scissile amide bond, thereby activating the carbonyl carbon for nucleophilic attack by water. This low-energy conformation, with the enzyme, water and substrate set up for intramembrane hydrolysis, overlaps remarkably well, particularly in the active site, with the new cryo-EM structure with TMD mimetic **SB-250** (Fig. 2e; Extended Data, Fig. 7). Thus, the new cryo-EM structure provides important support for the GaMD *in silico* simulations.

Given this structural confirmation, the six PSEN1 FAD mutations analyzed for comprehensive effects on γ-secretase cleavage of APP substrate were then examined for effects on activating the ε cleavage step using GaMD simulations. Consistent with the biochemical analysis, most of the six PSEN1 FAD mutations visited the active conformation less frequently, if at all.^30^ Analysis of GaMD simulations of the FAD-mutant E-S complexes further revealed that all six reduced conformational fluctuations compared with the WT E-S complex, in particular for PSEN1 TM6a and the associated region of APP substrate inserted into the active site (Fig. 2f; Extended Data, Fig. 8). The reduced conformational flexibility suggests a mechanism by which these mutations decrease ε cleavage, as enzyme catalysis is a dynamic process that requires conformational rearrangements. The reduced flexibility further suggests that the FAD-mutant E-S complexes are more stable: Because the substrate is enveloped by PSEN1 TMDs, dissociation of substrate from the enzyme should be more difficult. Additional GaMD simulations were performed for the Aβ49→Aβ46 trimming step. Remarkably, only P117L PSEN1 prevented the Aβ49/γ-secretase complex from visiting the active conformation (Fig. 2g), consistent with results from LC-MS/MS analysis of this trimming step (Fig. 1g). The complex of P117L PSEN1 γ-secretase with Aβ49 was also found to be the least flexible among the FAD mutations examined compared to the WT enzyme-Aβ49 complex (Extended Data, Fig. 9).

To test effects of FAD mutations on the stability of γ-secretase E-S complexes in cultured cells, we conducted fluorescence lifetime imaging microscopy (FLIM).^31^ WT or FAD-mutant PSEN1 was exogenously expressed together with C99 in HEK293 cells in which endogenous PSEN1 and PSEN2 were knocked out through CRISPR/Cas9 gene editing.^32^ The C99 construct contains human APP C99 with the N-terminal signal peptide sequence for membrane insertion and secretory pathway destination and a C-terminal near-infrared fluorescence protein: miRFP720 (C99-720). After fixing and permeabilization, primary antibodies were added that interact with the N-terminal region of C99/Aβ (mouse antibody 6E10) and to an epitope on γ-secretase component nicastrin that is proximal to the N-terminus of bound APP substrate in cryoEM structures (rabbit antibody NBP2-57365). Secondary antibodies conjugated to fluorophore were then added: anti-mouse IgG antibody conjugated to Alexa Fluor™ 488 and anti-rabbit IgG antibody conjugated to Cy3 (Fig. 3a). Although the 6E10 antibody reacts with both C99 and Aβ, dividing the 6E10-Alexa488 emission by that of C99-720 allows detection of cell compartment(s) with lower or higher 6E10 Alexa488/C99 720 ratios as sites where C99 or intracellular Aβ, respectively, are enriched (Fig. 3a).^33,34^ Selected cells (n = 7-10) were analyzed by FLIM, with the C99 or Aβ-rich regions of interest (ROIs; 136-141) quantified for each sample for average fluorescence lifetime of Alexa Fluor™ 488 (Fig. 3b). For each PSEN1 FAD-mutant sample, fluorescence lifetime was significantly reduced compared to that seen with WT PSEN1, both in the C99 and Aβ-rich areas, indicating that more C99/Aβ is proximal (i.e., bound) to FAD-mutant γ-secretase (Fig. 3c). Similar results were seen upon testing effects on FLIM using four FAD mutations located in the C99 TMD (Fig. 3d). Use of the acceptor antibody with an epitope on the other side of the protease complex (i.e., distal to the N-terminus of bound C99/Aβ) did not lead to any reduction in fluorescence lifetime (Extended Data, Fig. 10).

**Fig. 3.**
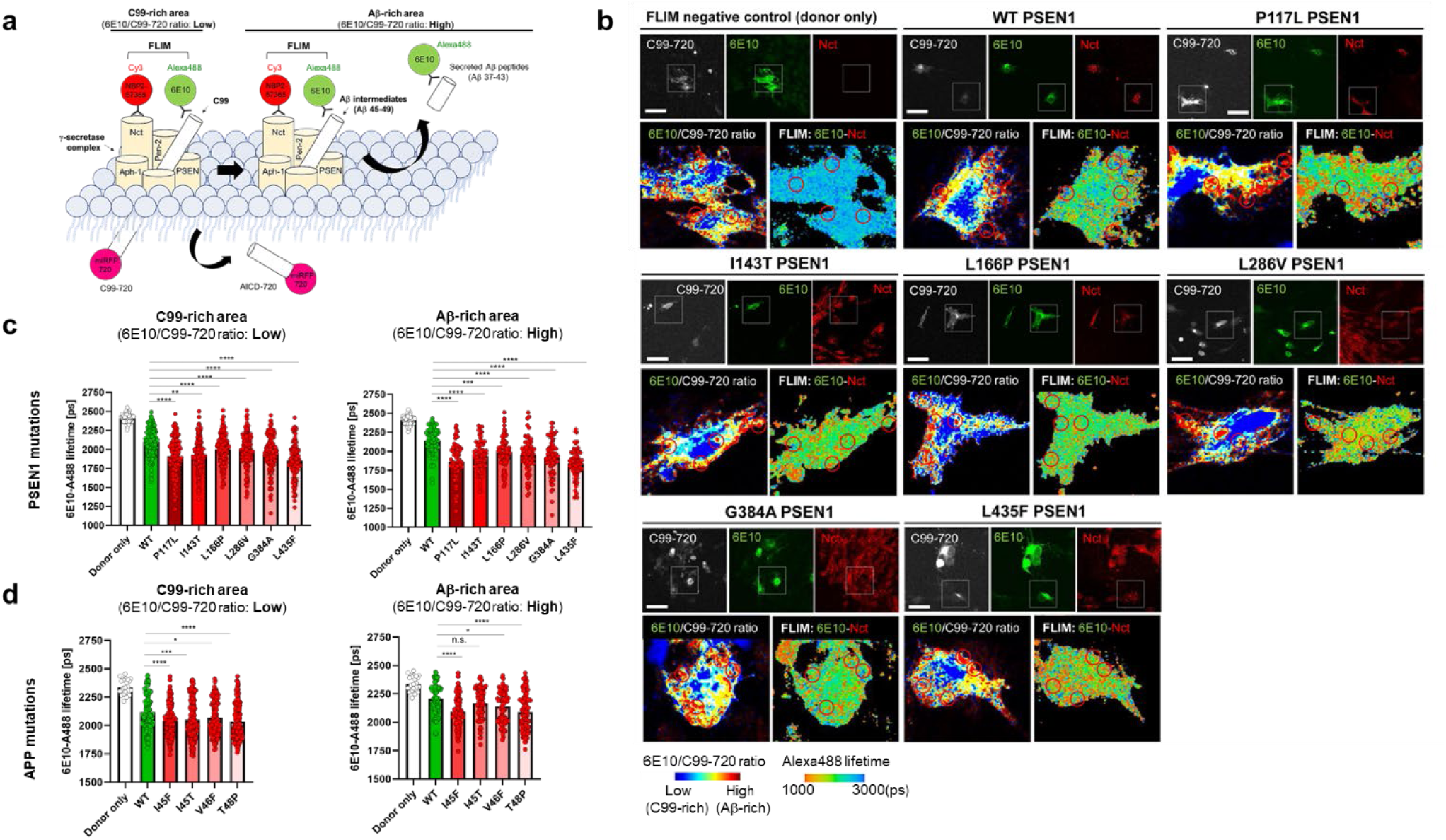
FAD PSEN1 mutations stabilize γ-secretase complexes bound to APP substrate and/or Aβ peptide intermediates. **a**, Design of fluorescence lifetime imaging microscopy (FLIM) experiments. **b**, Microscopic imaging with pseudo color analysis to identify C99 or Aβ intermediates-rich subcellular areas (regions of interest, ROIs; circles), followed by FLIM in HEK293 cells. **c**, Quantification of FLIM results reveal FAD mutations in PSEN1 and **d**, C99 increase stabilization of enzyme-C99 (left panels) or enzyme-Aβ intermediate complexes (right panels) in intact cells. For all graphs, n = 136-141 ROIs from 7-10 cells, one-way ANOVA and the Tukey’s multiple comparisons test, n.s., p>0.05* p<0.05, ** p<0.01. *** p<0.001, ****p<0.0001.

To develop a convenient *in vivo* system to probe pathogenic mechanisms in FAD, we generated transgenic *C. elegans* lines that coexpress C99 and human PSEN1, with and without FAD mutations, under control of the pan-neuronal *rgef-1* promoter (Fig. 4a). Parental line *juIs1* expresses synaptobrevin fused to GFP under control of the *unc-25* promoter, enabling visualization of GABAergic synaptic puncta along the dorsal and ventral nerve cords (Fig. 4a). Expression of C99 with the APP signal sequence allows membrane insertion, while expression of human PSEN1 leads to replacement of *C. elegans* orthologues sel-12 and hop-1 in endogenous γ-secretase complexes. Human PSEN1 can fully rescue *SEL-12* mutant phenotypes.^35,36^ Thus, C99 and functional PSEN1/γ-secretase complexes should be reconstituted into neuronal membranes of the roundworm for intramembrane proteolysis, allowing the testing of effects of FAD mutations. Coexpression of WT C99 and WT PSEN1 led to *C. elegans* lines with lifespans and numbers of synaptic puncta indistinguishable from those of the parental line (Extended Data Fig. 11 and Extended Data Table 2). In contrast, coexpression of I45F FAD-mutant C99 (Aβ numbering; Iberian APP mutation^37^) and WT PSEN1 gave transgenic lines with substantially reduced lifespans and age-dependent loss of synaptic puncta beginning on day 3-4 of adulthood (Fig. 4b-d; Extended Data Fig. 12; Extended Data Tables 3-6). These effects required coexpression of WT PSEN1, as monogenic lines expressing only I45F C99 displayed normal or near-normal lifespans and numbers of synaptic puncta. Thus, the neurodegenerative phenotype is due to the interaction of I45F C99 with WT PSEN1, with the latter likely incorporated as the catalytic component of the γ-secretase complex.

**Fig. 4.**
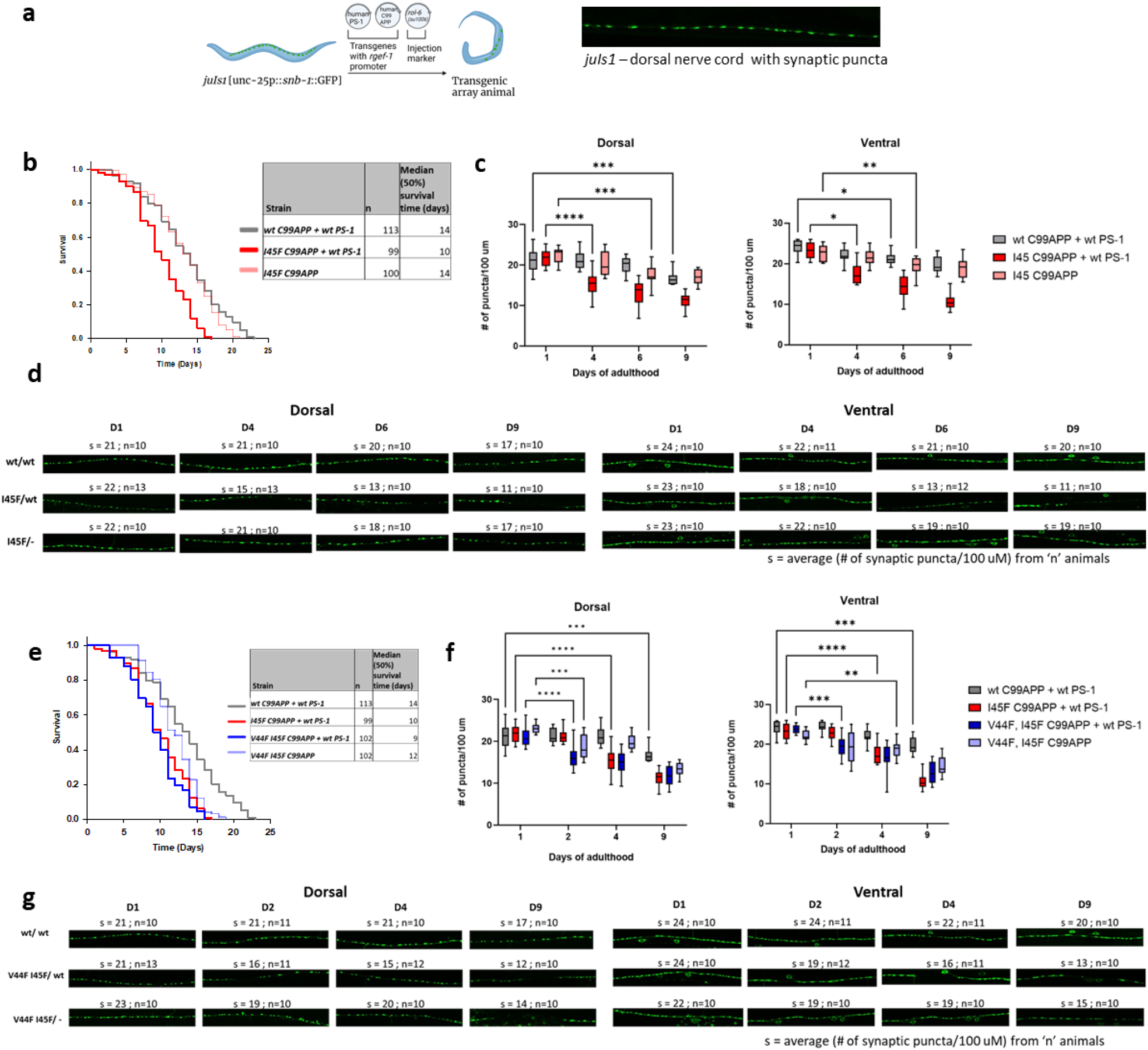
*C. elegans* model of FAD show reduced lifespan and age-dependent synaptic degeneration that is independent of Aβ42 production. **a**, Design of *C. elegans* transgenic model and visualization of synaptic puncta in the parental *juIs1* line. Transgenic lines were obtained by microinjection of human APP C99 and/or human PSEN1 (driven by pan-neuronal *rgef-1* promoter) into the parental line *juIs1. rol-6* (dominant roller) was used as the coinjection marker to select for transformants. *Right*: a representative confocal image showing a series of synaptic puncta marked by GFP-tagged synaptobrevin along the dorsal nerve cord of *juIs1*. **b,** Life span of double transgenic lines C99 + PSEN1 and C99 I45F + PSEN1 and monogenic line C99 I45F. Kaplan-Meier curves of n animals show the fraction of animals alive at different days. **c**, Quantification of the dorsal and ventral synaptic puncta in the transgenic lines. Average number of synaptic puncta per 100 μm in n transgenic worms for each day is shown in the vertical box plots. Horizontal lines at the top and bottom of each box represent the maximum and minimum values, respectively. Upper and lower ends of a box mark quartiles Q1 and Q3 values, respectively. The horizontal line inside the box shows the median value and marks Q2). Two-way ANOVA for all possible pairs using Tukey’s post hoc test, *p≤ 0.05, **p≤ 0.01, ***p≤ 0.001, ****p< 0.0001. Statistical differences shown only for earliest day significance was seen compared to Day 1 in a given line. **d**, 100 μm sections of representative confocal microscopic images dorsal (left) and ventral (right) nerve cords of transgenic animals. ‘s’ denotes average number of synaptic puncta per 100 μm from n animals. **e**, Life span of double transgenic line C99 V44F/I45F + PSEN1 and monogenic line C99 V44F/I45F compared with C99 + PSEN1 and C99 I45F + PSEN1 lines. **f**, Quantification of dorsal and ventral synaptic puncta in these transgenic lines. Two-way ANOVA for all possible pairs using Tukey’s post hoc test, *p≤ 0.05, **p≤ 0.01, ***p≤ 0.001, ****p< 0.0001. Statistical differences shown only for earliest day significance was seen compared to Day 1 in a given line. **g**, 100 μm sections of representative confocal microscopic images of dorsal and ventral synaptic puncta in these transgenic lines. The same data used in panels b-d for C99 + PSEN1 and C99 I45F + PSEN1 were reproduced in panels e-g for direct comparison with C99 V44F/I45F + PSEN1 and C99 V44F/I45F. All experiments shown in Fig. 4 were repeated using independent transgenic lines, with the results shown in Extended Data Figs. 12 and 14.

The I45F FAD APP mutation leads to a ∼15-to 30-fold increase in the Aβ42/Aβ40 ratio.^13,38,39^ This is due to near-complete block of the Aβ46→Aβ43 proteolytic step,^13^ because a phenylalanine residue in the P2’ position of substrate (i.e., two residues C-terminal to the cleavage site) is highly disfavored.^39^ To address whether the synaptotoxic effects of I45F C99 are attributable to Aβ42, we installed an additional V44F mutation, to block Aβ45→Aβ42 proteolysis. *In vitro* γ-secretase assays with LC-MS/MS analysis of small peptide coproducts confirmed that the V44F/I45F double mutation completed blocked both Aβ45→Aβ42 and Aβ46→Aβ43 trimming steps (Extended Data Fig. 13a-c). Moreover, generation of HEK293 cells stably expressing WT, I45F and V44F/I45F C99 revealed that both mutant forms of C99 comigrate with γ-secretase by native PAGE (Extended Data Fig. 13d-f), providing additional evidence for stalled E-S complexes in cells. V44F/I45F C99 generates essentially no Aβ42 (ref. 40; Extended Data Fig. 13); nevertheless, coexpression of this double mutant with WT PSEN1 in *C. elegans* did not rescue the phenotype seen with I45F C99 + WT PSEN1 (Fig. 4e-g; Extended Data Fig. 14; Extended Data Tables 3-6). Indeed, synaptic loss began even earlier, on day 2 of adulthood. A less severe phenotype was seen with the monogenic V44F/I45F C99 lines, without exogenous WT PSEN1 expression, possibly due to dysfunctional interaction of this double mutant with endogenous worm γ-secretase complexes. Nevertheless, the complete lack of any rescuing effect of V44F/I45F C99 in the presence of WT PSEN1 indicates that the neurodegenerative phenotype observed is independent of Aβ42.

To test whether the neurodegenerative phenotype is dependent on any form of Aβ, we coexpressed V50F/M51F C99 and WT PSEN1. This double mutation essentially blocks ε cleavage and Aβ production, yet V50F/M51F C99 can bind to γ-secretase and compete with other substrates (i.e., it forms stabilized E-S complexes with little or no processing to Aβ peptides).^39^ Coexpression with WT PSEN1 resulted in lines with similar reduced lifespan and number of synaptic puncta as seen with I45F and V44F/I45F C99 (Fig. 5a-c; Extended Data Fig. 15; Extended Data Tables 3-6). These findings suggest that stalled C99/γ-secretase E-S complexes can cause neurodegeneration in the absence of Aβ production. Intriguingly, while synaptic puncta were reduced quite early—as soon as Day 1 of adulthood—in the V50F/M51F C99 + WT PSEN1 lines, nerve cord synapses apparently regenerated by Day 7-9 to levels seen in the WT C99 + WT PSEN1 lines, suggesting compensatory mechanisms to counter synapse loss. We then tested the effects of PSEN1 FAD mutant L166P, which leads to >90% reduction in ε cleavage of C99 (Fig. 1b and ref. 41). Neuronal expression of L166P PSEN1 resulted in reduced lifespan with or without coexpression of WT C99 (Fig. 5d; Extended Data Fig. 16a; Extended Data Tables 3-6). Age-dependent loss of synaptic puncta was also seen in the L166P + WT C99 lines (Fig. 5e,f; Extended Data Fig. 16b,c); unfortunately, this analysis was not possible with L166P lines, due to their over-sensitivity (death) upon exposure to the anaesthetizing agent. We note, however, that in all the other *C. elegans* transgenic lines, synaptic loss always accompanies reduced lifespan. Thus, while FAD-mutant I45F C99 required coexpression of WT PSEN1 for a neurodegenerative phenotype, it is likely that FAD-mutant L166P PSEN1 does not require WT C99. These results suggest that stalled γ-secretase bound to other substrates besides C99 can trigger synaptic loss.

**Fig. 5.**
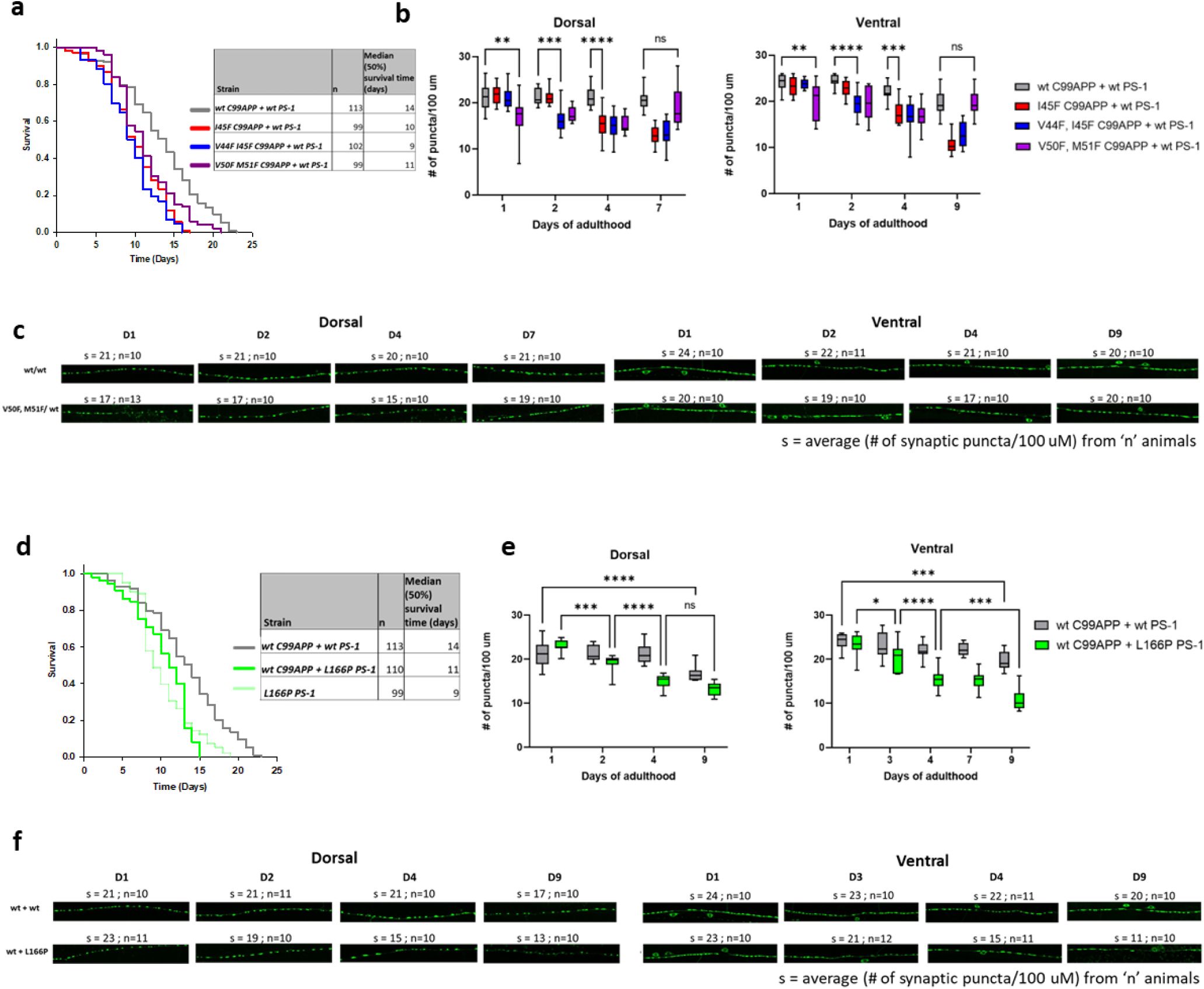
Neurodegeneration phenotype of FAD *C. elegans* models does not require Aβ production or C99. **a**, Life span analysis of double transgenic line C99 V50F/M51F + PSEN1 compared with C99 + PSEN1, C99 I45F + PSEN1 and C99 V44F/I45F + PSEN1 lines. **b**, Quantification of dorsal and ventral synaptic puncta in these transgenic lines. Two-way ANOVA for all possible pairs using Tukey’s post hoc test, *p≤ 0.05, **p≤ 0.01, ***p≤ 0.001, ****p< 0.0001. Statistical differences shown only for earliest day significance was seen between a given line versus C99 + PSEN1. **c**, 100 μm sections of representative confocal microscopic images of dorsal and ventral synaptic puncta in these transgenic lines. The same data used in Fig. 4 for C99 + PSEN1, C99 I45F + PSEN1 and C99 V44F/I45F + PSEN1 were reproduced in panels b-d for direct comparison with C99 V50F/M51F + PSEN1. **d**, Life span analysis of double transgenic line C99 + PSEN1 L166P and single transgenic line PS1 L166P compared with C99 + PSEN1. **e**, Quantification of dorsal and ventral synapses in double transgenic lines C99 + PSEN1 and C99 + PSEN1 L166P. *Note*: PS-1 L166P monogenic lines were sensitive to the anesthetic agent, phenoxy propanol, leading to dying on agarose pads during sample preparation for imaging. Thus, confocal imaging of these animals was not possible. Two-way ANOVA for all possible pairs using Tukey’s post hoc test, *p≤ 0.05, **p≤ 0.01, ***p≤ 0.001, ****p< 0.0001. Statistical differences shown only for days significance was seen within a given line. **f**, 100 μm sections of representative fluorescence microscopic images of dorsal and ventral synaptic puncta in these transgenic lines. The same data used in Fig. 4 for C99 + PSEN1 were reproduced in panels d-f for direct comparison with C99 + PSEN1 L166P and PSEN1 L166P (lifespan) or with only C99 + PSEN1 L166P (synaptic puncta).. All experiments shown in Fig. 5 were repeated with independent transgenic lines, with the results shown in Extended Data Figs. 15 and 16.

Difficulties in elucidating the pathogenic trigger(s) of Alzheimer’s disease and discovering effective therapeutics suggests that entities and processes beyond Aβ may be primarily responsible for initiating the cascade of events leading to neurodegeneration and dementia.^2,3^ Focusing on FAD should simplify identification of pathogenic mechanisms, as these rare variants of Alzheimer’s disease are caused by dominant missense mutations in the substrate and enzyme that produce Aβ. In the present study, full analysis of effects of FAD mutations on all the proteolytic steps in γ-secretase processing of C99 revealed that early events (ε cleavage and/or the first and second trimming steps) are commonly deficient. Elucidation of the structure of γ-secretase bound to a full TMD substrate-based probe provided a snapshot of the enzyme poised at the transition state of intramembrane proteolysis, and this structure matched a molecular dynamics model of the activated enzyme. The development of this *in silico* model allowed study of the effects of FAD mutations on the structural dynamic mechanism of γ-secretase processing of C99 substrate and Aβ49 intermediate, revealing that FAD-mutant enzyme-substrate (E-S) complexes that are proteolytically deficient occupy a smaller volume of conformational space. This reduced flexibility suggested that the mutant E-S complexes are stabilized while they are stalled in their proteolytic activities, and this idea was supported by reduced fluorescence lifetimes of labeled antibody probe combinations targeting E-S complexes. The development of a *C. elegans* model for FAD then provided a convenient *in vivo* system to interrogate pathogenic mechanisms, with results pointing to stalled, stabilized E-S complexes as the trigger of synaptic degeneration. These findings contradict an earlier report from Szaraga *et al.*, which suggested FAD mutations *de*stabilize E-S complexes, particularly those involving Aβ_n_ intermediates.^42^ However, that study did not test E-S complex stability *per se*. Here we provide a structural dynamic mechanism for stabilized FAD E-S complexes and FLIM results in intact cells to measure WT vs. FAD E-S complex stability.

The evidence presented here does not exclude roles for Aβ42 and its aggregated forms in FAD pathogenesis. The dysfunctional proteolytic processing caused by FAD mutations typically—but not always^43^—increases Aβ42/Aβ40, primarily through decreased Aβ40 production, thereby increasing the propensity of Aβ42 to aggregation. Such aggregation can lead to activation of glial cells and neuroinflammation that can aggravate the pathogenic process.^44^ Moreover, recently reported human trial results showed that an anti-Aβ antibody, lecanemab (Leqembi), cleared Aβ plaques from the brain and modestly slowed the rate of cognitive decline in Alzheimer’s disease.^8^ The modest effect on cognitive decline, however, suggests Aβ may not be a primary disease driver. We show here that the stalled process—not the products—of γ-secretase proteolysis of substrates can trigger age-dependent synaptic loss and reduce lifespan in *C. elegans*. In FAD, this stalled process can result from missense mutations in either substrate (APP) or enzyme (presenilin)(Fig. 6). Among the >100 identified substrates for γ-secretase, FAD mutations are only found in APP, and this may be due to its high expression in the brain^45^ along with its constitutive processing by β-secretase to C99.^46^

**Fig 6.**
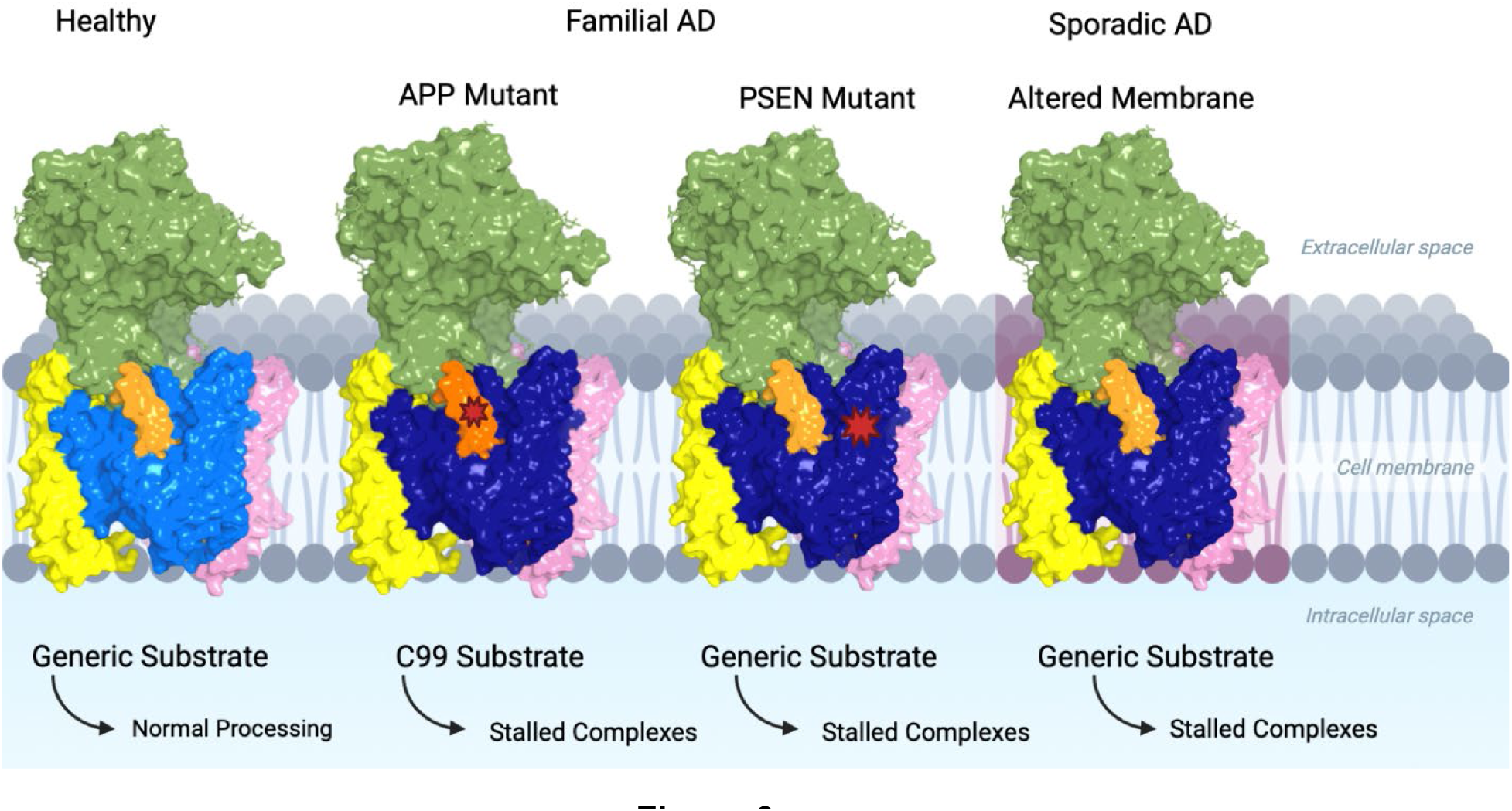
Model for how FAD mutations in APP and presenilins trigger pathogenesis and possible implications for sporadic Alzheimer’s disease. In healthy neurons, the γ-secretase complex can effectively process C99 and the enzyme’s many other membrane-bound substrates. FAD mutations in APP result in stalled γ-secretase complexes bound to C99 substrate or Aβ intermediate. FAD mutations in presenilin result in stalled interaction between γ-secretase and its substrates (not restricted to C99). In sporadic late-onset AD, in the absence of mutations in APP or presenilins, other factors, such as altered membrane fluidity, may stabilize E-S complexes, thereby triggering synaptic degeneration.

While our findings implicate stalled E-S complexes as triggers of FAD pathogenesis, they do not address whether stalled complexes do so through reduced proteolysis of critical γ-secretase substrates (in accord with the Presenilin Hypothesis^47^) or through a gain of toxic function of the stalled complexes *per se*. However, dominant mutations in PSEN1 that lead to loss of function through nonsense-mediated decay are associated with dominantly inherited skin disease and not neurodegeneration,^48^ suggesting stalled E-S complexes as such may trigger neurodegeneration. Interestingly, stalled ribosomes provide a precedent for slowed or stuck processive E-S complexes triggering gain of function (in this case, specific cellular stress responses).^49^ The question of gain versus loss of function mechanisms of synaptic degeneration will be important for future investigations, along with determining the downstream pathways and networks altered by stalled γ-secretase/substrate complexes. Regarding possible relevance of these findings to sporadic late-onset Alzheimer’s disease, we speculate that stalled γ-secretase bound up with substrate, perhaps as a consequence of altered membrane composition and properties (e.g., reduced fluidity) in the aging brain,^50^ may trigger this most common form of the disease (Fig. 6). Regardless, a key role of stabilized γ-secretase E-S complexes in FAD pathogenesis has implications for drug discovery for this rare genetic form of the disease, suggesting that the search for stimulators of the stalled complexes, to correct the dysfunctional proteolysis, would be worthwhile.

**Supplemental Material.** Methods, additional references; supplementary information, figures, and tables; acknowledgements; details of author contributions and competing interests; and statements of data availability are available in supplemental material.

