## Extended Data for "Alzheimer mutations stabilize synaptotoxic γ-secretase-substrate complexes"

### Supplemental Material for: Alzheimer mutations stabilize synaptotoxic γ-secretase-substrate complexes

#### Materials and Methods

##### $^{15}N$, $^{13}C$ labelled C100-FLAG expression and purification

*E. coli* BL21 cells were grown with shaking in minimal media with 20% $^{13}C$ glucose (Cambridge Isotope Laboratories) and $^{15}NH_4Cl$ (Cambridge Isotope Laboratories) at 37 $^0C$ until $OD_{600}$ reached 0.8. The minimal media composition is listed in Extended Data Fig 2. Cells were induced with 0.5 mM IPTG and grown for 3 h. Cells were harvested by centrifugation, resuspended in lysis buffer composed of 25 mM Tris pH 8 and 1% Triton X-100, and lysed passing through a French press three times. Cleared lysate was incubated with anti-FLAG M2-agarose beads (Sigma-Aldrich) for 16 h with shaking at 4 $^oC$. The beads were washed 3 times with lysis buffer, and C100-FLAG protein was eluted with buffer composed of 100 mM glycine at pH 2.7 and 0.25% NP-40, followed by neutralization with pH 8 Tris buffer and storage at -80 $^0C$. The identity and purity of C100-FLAG was analyzed by SDS/PAGE with Coomassie staining and MALDI-TOF mass spectrometry.

##### FAD-mutant γ-secretase constructs

Four monocistonic pMLINK vectors, pMLINK-PSEN1, pMLINK-Aph1 (with C-terminal HA epitope tag), pMLINK-NCT (with C-terminal V5 and 6XHIS epitope tags) and pMLINK-Pen-2 (with N-terminal STREP and FLAG epitope tags), were constructed as previously described.[1] FAD mutations were made in the PSEN1 coding region of pMLINK-PSEN1 by site-directed mutagenesis (QuikChange Lightning Multi-Site Directed Mutagenesis kit, Agilent). Each vector has LINK1 and LINK2 sequences flanking the gene of interest. LINK1 harbors a Pac1 restriction site and LINK2 harbors both PacI and Swa1 restriction sites. Mutated pMLINK-PSEN1 vector was treated with restriction enzyme Pac1 and electrophoresed through 1% agarose gel to separate and purify the FAD-mutant PSEN1 DNA. Similarly, pMLINK-APhH1 was treated with SwaI restriction enzyme to linearize the vector followed by electrophoresis through 1% agarose gel and gel band isolation. The PSEN1 fragment and linearized pMLINK-APhH1 vector were treated with T4 polymerase for 20 min at ambient temperature in the presence of dCTP or dGTP, respectively. The purified T4 polymerase-treated PSEN1 fragment was inserted into purified linearized pMLINK-Aph1 by ligation independent cloning (LIC) to create bicistronic pMLINK-Aph1-PSEN1 vector. Similarly, bicistonic pMLINK-Pen2-Nicastrin was created using the LIC method. Finally, the two bicistronic vectors were used to make the tetracistronic vector (pMLINK-PEN-2-nicastrin-APH-1-PS1) by the LIC method (See Extended Data Fig. 1 for illustration).

**γ-Secretase expression and purification**

γ-Secretase was expressed and purified from HEK 293F cells as described previously.[2-4] Briefly, HEK 293F cells were grown in unsupplemented Freestyle 293 media (Life Technologies, 12338-018) until cell density reached 2 x $10^6$ cells/mL. For transfection, media was replaced with fresh Freestyle 293 media. 150 mg pMLINK tetracistronic vector and 450 mg polyethylenimines of 25 kDa (PEI) was mixed in 5 mL of Freestyle media followed by incubation for 30 min at ambient temperature. The incubated DNA/PEI mixture was added to the cell culture, which after 24 h was sub-cultured in three flasks and further grown for 36 h before harvesting. Cells were pelleted by centrifugation and resuspended in buffer consisting of 50 mM MES pH 6.0, 150 mM NaCl, 5 mM $CaCl_2$ and 5 mM $MgCl_2$, and lysed by French press. Unbroken cells and debris were pelleted by centrifugation at 3000 x g for 10 minutes. The supernatant was ultracentrifuged at 100,000 x g for 1 h to isolate the membrane pellet. The membrane pellet was washed with 0.1 M sodium bicarbonate pH 11.3 by successive passage through syringes with 18-, 22-, 25- and 27-gauge needles, followed by ultracentrifugation at 100,000 x g for 1 h. The pellet was resuspended in 50 mM HEPES pH 7, 150 mM NaCl and 1% CHAPSO, and incubated in ice for 1 h, followed by ultracentrifugation at 100,000 x g. The supernatant was mixed with anti-FLAG M2-agarose beads (Sigma-Aldrich) and TBS with 0.1% digitonin and incubated for 16 h at 4 $^{o}$C. Beads were washed three times with TBS/0.1% digitonin before eluting the γ-secretase complex with buffer consisting of 0.2 mg/mL FLAG peptide in TBS/ 0.1% digitonin. The eluate was stored at -80 $^{0}$C until further use.

**In vitro γ-secretase assay**

In vitro γ-secretase assay was performed as described previously.[5] Briefly, 30 nM of WT or FAD mutant γ-secretase was preincubated for 30 min at 37 °C in assay buffer composed of 50 mM HEPES pH 7.0, 150 mM NaCl, and 0.25% 3-[(3-cholamidopropyl) dimethylammonio]-2-hydroxy-1-propanesulfonate (CHAPSO), 0.1% phosphatidylcholine and 0.025% phosphatidylethanolamine. Reactions were started by adding either the light- or heavy-isotope form of C100-FLAG substrate (3 μM final concentration) and incubating at 37 $^{o}$C for 16 h. The reactions were stopped by flash freezing in liquid nitrogen and stored at -20 $^{o}$C.

**Tri- and tetra peptide products analysis by LC-MS/MS**

Small peptides were analyzed using an ESI Quadrupole Time-of-Flight (Q-TOF) mass spectrometer (Q-TOF Premier, Waters) by LC-MS/MS as described previously.[5] The reaction mixture consisting of light C100-FLAG/WT γ-secretase and reaction mixture consisting of heavy C100-Flag/FAD γ-secretase were mixed 1:1 prior to injection onto a C18 analytical chromatography column and eluted with a step gradient of 0.08% aqueous formic acid, acetonitrile, isopropanol, and a 1:1 acetone/dioxane mixture. The three most abundant fragments from collision-induced dissociation were identified by tandem MS for each small peptide. Similarly, various concentrations of the synthetic peptides' standards (>98% purity,

New England Peptide) were dissolved in assay buffer and loaded onto a C18 analytical chromatography column for LC-MS/MS analysis. To obtain a peptide chromatographic area, the signals from the 3 most abundant ions were summed using an ion mass width of 0.02 unit. Data were acquired in “V” mode.

**Immunoblotting of AICD products**

For quantitative immunoblotting of total AICD-Flag products, γ-secretase reaction samples and C100-FLAG standards were run on SDS-PAGE 4-12% Bis-Tris gels and transferred to PVDF membranes. Membranes were blocked with 5% dry milk for 1 h at ambient temperature and treated with anti-Flag M2 antibodies (Sigma-Aldrich) for 16 h at 4 $^{o}$C. The blot was then washed 3 times and incubated with anti-mouse secondary antibodies for 1 h at ambient temperature. The membrane was washed and imaged for chemiluminescence, and bands were analyzed by densitometry. A standard curve established from a concentration range of C100-FLAG protein was used to calculate AICD-FLAG product concentration from the enzyme reaction mixtures.

**Detection of AICD species by mass spectrometry**

AICD-FLAG in the reaction mixture was immunoprecipitated with anti-FLAG M2 beads (Sigma-Aldrich) in 10 mM MES pH 6.5, 10 mM NaCl, 0.05% DDM detergent for 16 hours at 4 $^{o}$C. AICD products were eluted from the anti-FLAG beads with acetonitrile:water (1:1) with 0.1% trifluoroacetic acid. The elutes were analyzed on a Bruker Autoflex MALDI-TOF mass spectrometer.

**Quantification of Aβ40 and Aβ42 by ELISA**

To measure secreted Aβ peptides, conditioned media were collected after 48 h from confluent HEK293 cells (nontransfected control line and those stably transfected with variants of C99). Levels of secreted Aβ40 and Aβ42 in the conditioned media were measured with specific ELISA kits from Invitrogen according to the manufacturer's instructions. All samples were measured in triplicate. Similarly, levels of Aβ40 and Aβ42 derived from *in vitro* cleavage assays with purified γ-secretase complexes and C100Flag substrate were determined.

**Cell culture and DNA transfection for C99-expressing HEK293 cells**

Human embryonic kidney 293 (HEK293) cells stably expressing human C99, and its mutants were cultured in Dulbecco's modified Eagle's medium (DMEM; GIBCO, USA) at 37°C and 5% $CO_2$. All cell lines were grown in this medium supplemented with 10% fetal bovine serum (FBS; GIBCO, USA). HEK293 cells were transfected using Lipofectamine 3000 (Thermo Fisher Scientific, USA) with the cDNA of human C99 with the APP signal sequence inserted into pCMVvector containing a puromycin resistance gene (gift of L. Liu, Brigham and Women's Hospital, Boston). Two different mutant forms of C99 were transfected separately: 1) "C99-F", containing the I45F Iberian FAD mutation; 2) "C99-FF", further containing an artificial V44F mutation that blocks Aβ42 production. After transfection, cells were treated with puromycin (1 mg/ml) to select for cells stably expressing the C99 variants.

**Antibodies and Western blot analysis for C99-expressing HEK293 cells**

The following antibodies were used: Anti-GAPDH (Cell Signaling Technology, 97166T), Anti-PS1-NTF (Bio-Legend, 82340), anti-Aβ 6E10 (Bio-Legend, 803001), Anti-total Aβ (Cell

Signaling Technology, 8243T). Western blot analysis was performed using standard techniques. Cells were lysed in 1% digitonin and a protease inhibitor mixture, protein levels determined via BCA assay (Thermo Fisher Scientific, USA), and protein-normalized lysates were subjected to SDS-PAGE and transferred to PVDF membranes. Immunoblot analysis was then performed and visualized by the enhanced chemiluminescence method.

**Blue native–polyacrylamide gel electrophoresis (BN-PAGE) from C99-expressing HEK cells.** Cells were lysed in a native sample buffer (Thermo Fisher Scientific, USA) containing 1% digitonin and a protease inhibitor mixture. After centrifugation at 20,000 X g at 4 °C for 30 min, the supernatant was separated on a 3–12% BisTris gel (Thermo Fisher Scientific, USA) according to the instructions of the Novex BisTris gel system (Thermo Fisher Scientific, USA). The transferred blot was incubated in 20 mL of 8% acetic acid for 15 minutes to fix the proteins, rinsed with deionized water and analyzed with Western blotting.

**Sample preparation for cryo-EM study**

The γ-secretase complex was purified as previously described.[1] The peak fractions from gel filtration were concentrated to ~40 μM to incubate with 1.5 mM of transmembrane substrate mimitic SB-250 (final concentration) for 1 h at 4 °C. Cryo-EM grids were prepared with a Vitrobot (FEI). Aliquots of 4 μL of the SB-250-protein mixture were added to glow-discharged grids (Quantifoil Au R1.2/1.3). The grids were blotted for 3 s and frozen by liquid ethane, and then transferred to liquid nitrogen for storage.

**Data collection and processing for cryo-EM analysis**

The prepared grids were imaged on an FEI 300 kV Titan Krios electron microscope equipped with GIF Quantum energy filter (slit width 20 eV) and Gatan K3 Summit detector with a nominal magnification of 81,000 × (pixel size 1.0825). 8,324 micrographs were collected in total. MotionCor2 and Gctf were subsequentially used for motion correction and defocus value estimation, respectively. 3,556,137 particles were auto-picked with RELION (version 3.0) and were subjected to two-dimensional (2D) classification. 2, 182, 400 particles were then selected for 50 iterations of global angular search three-dimensional (3D) classification with a class number of 1 and step size of 7.5°. For the last six iterations (No. 45-50) of the global search, the local angular search 3D classification was performed with a class number of four, a step size of 3.75°, and a local search range of 15°. For the last iteration of the local search, particles from the good classes were merged and duplicated particles were removed, yielding 1,561,706 particles. Then we generated multi-reference models from one of the last iterations of local search 3D classification. The merged particles were applied to multi-reference-based 3D classification. Finally, 349,532 particles from the good classes were auto-refined, resulting in 3.4-Å reconstruction. After mask application, increasing box size and postprocessing, a final reconstruction of 2.6-Å was achieved based on the Fourier shell correlation (FSC) 0.143 criterion. Local resolution estimation was performed using RELION-3.0.

**Aβ49→Aβ46 molecular dynamics simulation system setup**

Starting from the "Active" WT conformation obtained from a previous study,[6] the amide bond between APP residue L49 and V50 was cleaved to prepare the starting structure. The PS1 FAD mutations, including G384A, I143T, L166P, L286V, L435F, and P117L, were computationally generated using the *Mutation* function of CHARMM-GUI.[7-10] Here, residue D385 in PS1 was protonated and the C-terminal of Aβ49 and N-terminal of AICD50-83 were charged to simulate γ-secretase activation for ζ cleavage of Aβ49 based on previous studies.[6,11] Other chain termini were capped with neutral patches (acetyl and methylamide).

The γ-secretase complexes were embedded in POPC membrane lipid bilayers and solvated in 0.15 M NaCl solutions using the CHARMM-GUI webserver.[7-10]

**Simulation protocols**

Gaussian-accelerated molecular dynamics (GaMD) simulations of WT and PS1 FAD mutant γ-secretase bound by APP-C83 (i.e., to simulate activation for ε cleavage) were obtained from a previous study[6]. The CHARMM36m force field parameter set[12] was used for the protein lipids. The simulation systems were initially energetically minimized for 5000 steps using the steepest-descent algorithm and equilibrated with the constant number, volume, and temperature (NVT) ensemble at 310 °K. They were further equilibrated for 375 ps at 310 °K with the constant number, pressure, and temperature (NPT) ensemble. Short conventional molecular dynamics (cMD) simulations were then performed for 10 ns using the NPT ensemble with constant surface tension at 1 atm and 310 °K. All-atom Peptide Gaussian accelerated molecular dynamics (Pep-GaMD) implemented in the GPU version of AMBER 20[13,14] was applied to simulate the effects of PS1 FAD mutations on γ-secretase activation for cleavage of Aβ49 to Aβ46. Selective boost potential was applied to the essential potential energy of the bound peptides (Aβ49 and AICD50-83). The threshold energy $E$ for adding total boost potential was set to the upper bound, i.e., $E_P = V_{minP} + (V_{maxP} – V_{minP}) / k_{0P}$, whereas the threshold energy $E_D$ for adding dihedral boost potential was set to the lower bound, i.e., $E_D = V_{maxD}$.[14,15] The upper limits of the boost potential standard deviations, $\sigma_{0P}$ and $\sigma_{0D}$, were set to 8.0 kcal/mol and 6.0 kcal/mol, respectively. The PepGaMD simulations involved an initial short cMD of 15 ns to calculate acceleration parameters and equilibration of added boost potentials for 60 ns. Three 600 ns independent production simulations with randomized initial atomic velocities were performed on the γ-secretase complexes.

**Simulation analysis**

The simulation trajectories were analyzed using VMD[16] and CPPTRAJ.[17] Root-mean-square fluctuations (RMSFs) of PS1 and substrate within the γ-secretase complexes were calculated by averaging the RMSFs calculated from individual PepGaMD simulations of each γ-secretase system. Changes in the RMSFs (ΔRMSF) from the WT to PS1 FAD mutant γ-secretase were calculated by subtracting the RMSFs of the WT from those of PS1 FAD mutant γ-secretase (Fig. 2f and Extended Data Fig. 7). The distance between Cγ atoms of catalytic aspartates PS1-D257 and D385 and distance between PS1 residue D385 (atom OD2) and Aβ49 residue V46 (atom O) were calculated. The PyReweighting[18] toolkit was applied for free energy calculations from the D257-D385 and D385-V46 distances for each system (Fig. 2g). A bin size of 1 Å and cutoff 500 frames in each bin was used to calculate the two-dimension (2D) potential mean force (PMF) free energy profiles.

**Plasmid DNA for Fluorescence Lifetime Imaging Microscopy Experiments**

The C99-720 plasmid, which contains APP signal peptide, human APP C99, FLAG tag, and miRFP720, was sub-cloned from the C99 miRFP720-miRFP670 (C99 720-670) biosensor[19] into pcDNA3.1 (+) empty vector using NheI/EcoRI. Then, mutagenesis was performed to introduce the stop codon after the miRFP720 sequence using primers FW: ATCGGCGTGATGGAAGAGTAAGAATTCTGCAGATATCCA and RV: TGGATATCTGCAGAATTCTTACTCTTCCATCACGCCGAT. The plasmid sequence was verified by the MGH CCIB DNA core.

**Immunocytochemistry**

PSEN1/2 dKO HEK293 cells[20] co-transfected with C99-720 and WT or FAD-mutant PSEN1 were fixed with 4% paraformaldehyde (PFA) (VWR, Radnor, PA), and permeabilized by 0.1% Triton-X100 (Sigma-Aldrich, St. Louis, MO). The permeabilized cells were then incubated with mouse monoclonal 6E10 (BioLegend, San Diego, CA) and rabbit polyclonal nicastrin antibodies (Novus Biologicals, LLC, Centennial, CO), followed by an Alexa Fluor™ 488 (FRET donor) or Cy3 (acceptor)-conjugated anti-mouse and rabbit IgG secondary antibodies, respectively (Thermo Fisher Scientific). For control experiment in Extended Data Fig. 10, primary antibody to PSEN1 Loop (EP2000Y; Abcam) was used in place of anti-nicastrin antibody. The slide was covered by a coverslip using Fluoromount-G™ Mounting Medium (Thermo Fisher Scientific) and stored at 4 ºC.

**Confocal microscopy and fluorescence lifetime imaging microscopy (FLIM)**

An Olympus FV3000 Confocal Laser Scanning Microscope (Tokyo, Japan) was used to perform confocal microscopy and FLIM. Lasers at 488 nm, 561 nm, and 640 nm were used to excite an Alexa Fluor™ 488, Cy3, and C99-720, and the emitted fluorescence was detected within 500-530 nm (6E10-Alexa 488), 560-590 nm (nicastrin-Cy3), and 700-800 nm (C99-720) using a 40x/0.95NA objective. Pseudo-colored images corresponding to the ratios of 6E10-Alexa Fluor™ 488 over C99-720 emission were generated in MATLAB (MathWorks, Natick, MA).

In FLIM analysis, a mode-locked Chameleon Ti: Sapphire laser (Coherent Inc., Santa Clara, CA) set at 850 nm was used to excite an Alexa Fluor™ 488 fluorophore (FRET donor), and the emission was collected using the ET525/50m-2p filter (Chroma Technology Corp, Bellows Falls, VT). The donor Alexa Fluor™ 488 lifetime was recorded using a high-speed photomultiplier tube (MCP R3809; Hamamatsu photonics, Hamamatsu City, Japan) and a time-correlated single-photon counting acquisition board (SPC-830; Becker & Hickl GmbH, Berlin, Germany). The acquired FLIM data were analyzed using SPC Image software (Becker & Hickl GmbH). The donor Alexa 488's average lifetime (t1) was first measured in the absence of the acceptor fluorophore (Cy3) (donor only/FLIM negative control). Then, the donor Alexa 488's average lifetime was recorded in the presence of the acceptor Cy3 fluorophore (t2). The proximity between the donor and acceptor (less than 5–10 nm apart) results in energy transfer from the donor to the acceptor (FRET present), yielding characteristic shortening of t2.

##### *C. elegans* transgenes

*C. elegans* was maintained on nematode growth medium (NGM) plates and fed *E. coli* OP50.[21] *juIs1* strain [unc-25p::snb-1::GFP + lin-15(+)] expresses GFP fused to synaptobrevin in presynaptic terminals of GABAergic dorsal and ventral nerve cord neurons.[22] Human APP C99 and PS-1 cDNA were synthesized by GeneArt through Thermo-Fisher. Human C99 constructs include DNA encoding APP signal peptide ($APP_{1\text{-}17}$: MLPGLALLLLAAWTARA) followed by APP C99. These constructs were designed to include worm introns. All human constructs were cloned in worm expression vector pEVL415 (P*rgef-1*:*htau40::gfp::unc-54* 3′UTR). The vector was linearized, with excision of *htau40:gfp* DNA, by restriction digestion with BamH1 and NgoM4. PCR was performed to amplify the genes of interest from the constructs carrying them using primers that were designed to have ~ 15 bp 5' extensions complementary to the 3' sticky ends of the linearized vector. Cloning was performed by In-Fusion cloning (TakaraBio) following manufacturer's protocol. *C. elegans* carrying extra-chromosomal arrays of transgenic plasmids (human C99APP and/or human PS-1) were generated using pRF4 as a co-injection marker.

Plasmids generated in the study :

1 pEVL545 [*Prgef-1*::signal peptide : human wt C99APP :: *unc-54* 3'UTR]

2 pEVL546 [*Prgef-1*::signal peptide : human C99APP (I45F) :: *unc-5*4 3'UTR

3 pEVL547 [*Prgef-1*::signal peptide : human wt PS-1 :: *unc-54* 3'UTR]

4 pEVL548 [*Prgef-1*::signal peptide : human L166P PS-1 :: *unc-54* 3'UTR]

5 pEVL549 [*Prgef-1*::signal peptide : human V44F I45F C99APP :: *unc-54* 3'UTR]

6 pEVL554 [*Prgef-1*::signal peptide : human V50F M51F C99APP :: *unc-54* 3'UTR]

Full sequences of the transgenes are provided as a supplementary file.

**_C. elegans_ maintenance and transgenesis**

Animals were maintained on nematode growth medium (NGM) plates and fed *E. coli* OP50.[21] Transgenic lines were obtained by injecting human constructs with co-injection marker into gonads of *juIs1* [*Punc-25*::SNB-1::GFP]. Specifically, a mix of human APP variant (8-10 ng/μl) and/ or human PS-1 variant (8-10 ng/μl) plus marker *rol-6* (60.8 ng/μl) was injected into *juIs1* animals, subsequently selecting for animals displaying the roller phenotype.

Strains generated in this study:

| | |
|---|---|
| wt C99APP + wt PS-1 1st line | *lhEx661* |
| wt C99APP + wt PS-1 2nd line | *lhEx662* |
| I45F C99APP + wt PS-1 1st line | *lhEx655* |
| I45F C99APP + wt PS-1 2nd line | *lhEx656* |
| I45F C99APP 1st line | *lhEx648* |
| I45F C99APP 2nd line | *lhEx649* |
| V44F I45F C99APP + wt PS-1 1st line | *lhEx650* |
| V44F I45F C99APP + wt PS-1 2nd line | *lhEx651* |

| | |
|---|---|
| V44F I45F C99APP 1st line | *lhEx652* |
| V44F I45F C99APP 2nd line | *lhEx653* |
| V50F M51F C99APP + wt PS-1 1st line | *lhEx663* |
| V50F M51F C99APP + wt PS-1 2nd line | *lhEx664* |
| wt C99APP + L166P PS-1 1st line | *lhEx657* |
| wt C99APP + L166P PS-1 2nd line | *lhEx658* |
| L166P PS-1 1st line | *lhEx659* |
| L166P PS-1 2nd line | *lhEx660* |

***C. elegans* lifespan analysis**

Lifespan experiments were performed by selecting L4 animals from a plate that contains mixed stage animals. Animals were transferred to fresh plates whenever necessary to avoid contamination with animals of successive generations. Animals were maintained at 20 ℃ and were checked for survival at least once in two days. Animals that did not move when prodded were considered dead. Survival curves, calculation of median lifespan and statistical analysis were performed in 'Sigmaplot' using Kaplan Meier (log-rank test) method.

**Microscopy and image analysis**

L4 animals were selected and were maintained at 20 ℃. Adult worms from day 1 to day 7 and day 9 were scored for the number of SNB-1::GFP puncta in the ventral and dorsal nerve cords. Adult animals were anesthetized using 0.5% 2-phenoxypropanol in M9 and mounted on 2% agarose pads. Imaging was done using Olympus FV1000 laser scanning confocal microscope at 60X magnification NA 1.42. Image settings and acquisition parameters were optimized using Fluoview histogram function. All images were subsequently acquired using identical parameters. Specific acquisition parameters for each image are saved in the Olympus Image Format metadata.

*C. elegans* were also imaged in the Microscopy and Analytical Imaging Resource Core Laboratory (RRID:SCR_021801) at The University of Kansas using a TCS SPE Laser Scanning Confocal Upright Microscope (Leica Microsystems, DM6-Q model), with the 488 nm laser line, a Leica 63X/1.3NA ACS APO oil objective, 12-bit spectral PMT detector and a Leica LAS X Imaging software (version 3.5.7.23225). Synaptic puncta signal were detected using 488 nm excitation, 500-520 nm emission range. Images were captured at 1024 x 1024-pixel resolution, no bidirectional scanning, and a zoom factor at 1.0.

Confocal images were processed by using Fiji (ImageJ) software [23]. Fluorescent confocal images were opened with Fiji, and the series of slices for each image were stacked by 'Z- project'. Each stacked image was then converted to binary image [24]. A threshold value is then identified for each image to resolve individual synaptic punctum, also ensuring all puncta are included in the range by noticing the correspondence with visually evident puncta. While applying the threshold for ventral cord, cell bodies or any nonspecific background fluorescent objects are excluded from the analysis. Particles within the size range of 0.2 - ꝏ (micron^2) were included in the analysis, excluding any that were on the edges along both axes using "Analyze Particle" command in ImageJ. As a result, the location (x,y coordinates) of every puncta was obtained automatically. These values were exported to MS-Excel. Average number of synaptic puncta per 100 µm is determined by calculating the distance between two adjacent puncta along the x-axis. Graphs were plotted using GraphPad Prism 9 software. ANOVA was performed in this software, and significance for all possible pairwise comparisons were calculated using a Tukey's multiple comparisons test. For all statistical measurements, a threshold of adjusted P value <0.05 was set to determine significance in multiple comparison tests. (ns for $P > 0.05$, * for $P \leq 0.05$, ** for $P \leq 0.01$, *** for $P \leq 0.001$, **** for $P \leq 0.0001$).

**Acknowledgments.** We thank L. Liu (Harvard Medical School/Brigham and Women's Hospital) for HEK293 cells with PSEN1/2 doubly knocked out through genome editing, P. Arafi (U. Kansas undergraduate researcher) for rendering Figure 6, E. Lundquist (U. Kansas) for access to injection apparatus for transgene insertion into *C. elegans*, and E. Rosa-Molinar and N. Martinez-Rivera for assisting with microscopy at the Microscopy and Analytical Imaging Research Resource Core Laboratory at U. Kansas. This work was supported by grants GM122894, AG66986 and AG79569 from the U.S. National Institutes of Health and a Pilot Project Grant from the University of Kansas Alzheimer Disease Research Center via NIH grant P30 AG072973 (M.S.W.); the National Natural Science Foundation of China (Project 81920108015), the National Key R&D Program (2020YFA0509300) from the Ministry of Science and Technology of China, and the Key R&D Program of Zhejiang Province (2020C04001), Frontier Research Center for Biological Structure, and Start-up funds from Westlake University (Y.S.); and grant 2121063 from the U.S. National Science Foundation (Y.M). This work used supercomputing resources with allocation award TG-MCB180049 through ACCESS, project M2874 through NERSC, the BigJay (NSF award MRI-2117449) and Research Computing Clusters at the University of Kansas.

**Database availability.** CryoEM map for the structure of γ-secretase bound to probe SB-250 has been deposited in the Electron Microscopy Data Bank (EMDB) under the ID code EMD-36948. Atomic model for the structure of γ-secretase bound to probe SB-250 has been deposited in the Protein Database (PDB) under the ID code 8K8E.

**Author contributions.** Conceived and designed experiments: S.D., R.Z., V.N., M.M., H.D., A.N., J.T.D., Y.M., B.D.A., Y.S., M.S.W. Performed experiments: S.D., R.Z., V.N., M.M., H.D., C.O., S.B., A.S., A.N., J.T.D. Analysis of data: S.D., R.Z., V.N., M.M., H.D., A.N., C.O., J.T.D., A.S., Y.M., B.D.A., Y.S., M.S.W. Contributed to the writing and editing of the manuscript: S.D., R.Z., V.N., M.M., H.D., J.T.D., Y.M., B.D.A., Y.S., M.S.W.

**Competing interests.** The authors have no competing interests to declare.

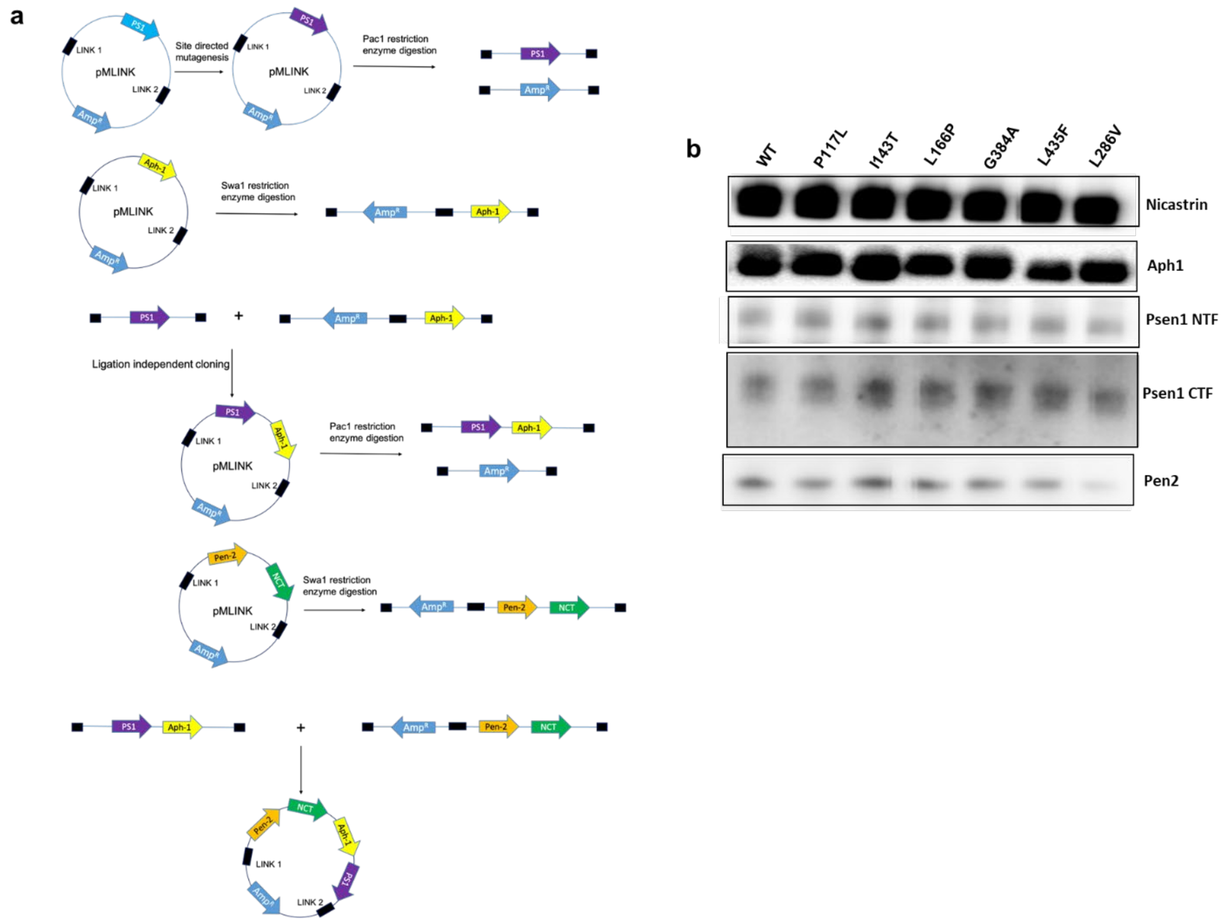

**Extended Data Figure 1. Generation and expression of pMLINK tetracistronic construct for FAD-mutant γ-secretase complexes. a,** FAD mutations were made in the PSEN1 coding region of pMLINK-PSEN1 by site-directed mutagenesis. FAD-mutant PSEN1 DNA was isolated after cleavage of Pac1 restriction sites in LINK1 and LINK2 regions. pMLINK-APhH1 was linearized via cleavage of a SwaI restriction site in LINK2. The PSEN1 fragment and linearized pMLINK-APhH1 were treated with T4 polymerase in the presence of dCTP or dGTP, respectively and combined by ligation independent cloning to create bicistronic pMLINK-Aph1-PSEN1 vector. Similarly, bicistonic pMLINK-Pen2-Nicastrin was created. Finally, the two bicistronic vectors were used to make the tetracistronic vector (pMLINK-PEN-2-nicastrin-APH-1-PS1). **b,** Western blot of all components of expressed and purified WT and FAD-mutant γ-secretase complexes, normalized for protein concentration.

| | |
|---|---|
| **Minimal media (500 ml)** | |
| $N_2HPO_4$ (anhydrous) | 3.4 g |
| $KH_2PO_4$ | 8.794 g |
| NaCl | 0.25 g |
| $^{15}$NH4Cl | 0.5 g |
| 20% $^{13}$C glucose | 10 ml |
| 1M $MgSO_4$.7H2O | 1 ml |
| 1M $CaCl_2$.2H2O | 10 μl |
| 0.5% Thiamine HCl 500 μl | |
| BME vitamins (Sigma) | 5 ml |
| 1MFeSO4.7H2O | 10 μl |

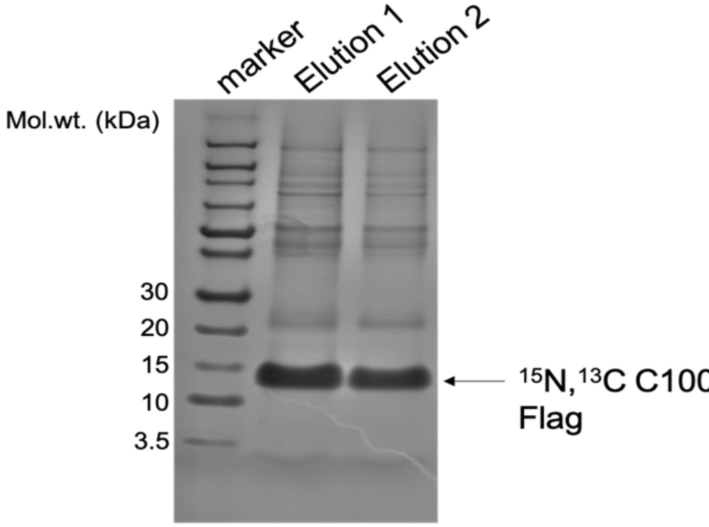

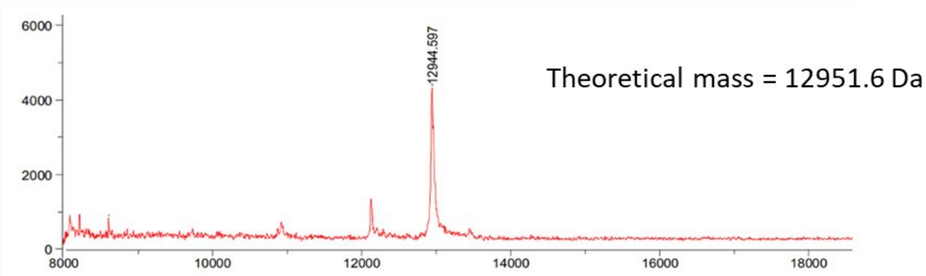

**Extended Data Figure 2. Production, purification and characterization of heavy isotopic C100Flag protein.** *E. coli* BL21 cells were grown in minimal media with 20% $^{13}$C glucose and $^{15}NH_4Cl$ (complete composition shown) and induced with 0.5 mM IPTG. Cell lysate was incubated with anti-FLAG M2-agarose beads, and C100-FLAG protein was eluted with 100 mM glycine buffer at pH 2.7 and 0.25% NP-40, followed by neutralization with pH 8 Tris buffer. The identity and purity of C100-FLAG was analyzed by SDS/PAGE with Coomassie staining and MALDI-TOF mass spectrometry. Note: Theoretical monotopic mass of light C100-FLAG is 12,265.05.

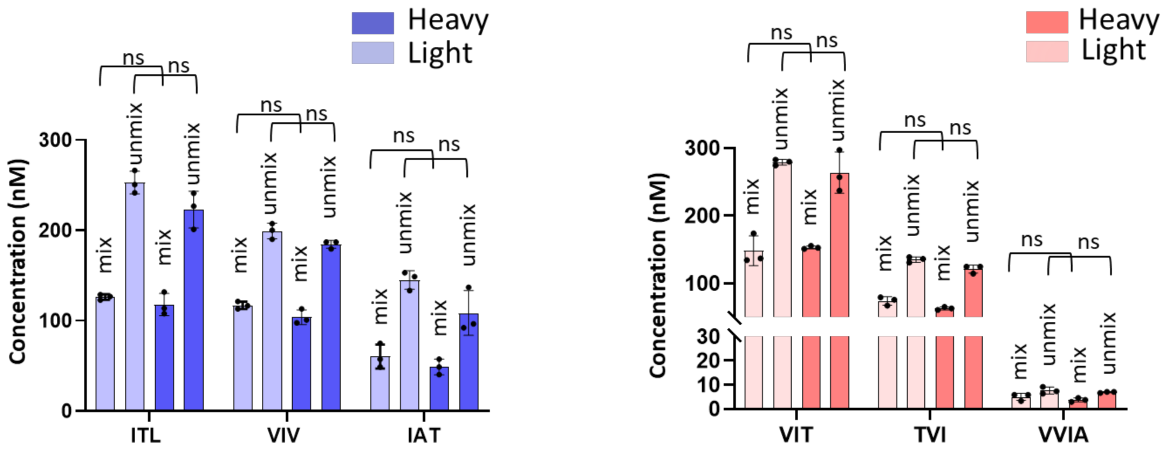

**Extended Data Figure 3. Validation of light- vs. heavy-isotopic substrate labeling method using WT γ-secretase.** Light- and heavy-isotope labeled C100Flag substrate were separately incubated with WT γ-secretase. After quenching, the two enzyme reactions were then mixed 1:1 for analysis of small peptide co-products of each proteolytic event. Shown are the concentrations of each co-product for both the unmixed and mixed samples. Note that light- and heavy-isotope labeled substrate gives equimolar levels of each small peptide, both in the unmixed and mixed samples. Mixed samples contain half the "light" or "heavy" co-product seen in the unmixed samples due to the 1:1 dilution. N=3, unpaired two-tailed T-test

**a**

[Aβ49] = [AICD 50-99] – [ITL]
[Aβ46] = [ITL] – [VIV]
[Aβ43 ]= [VIV] – [IAT]
[Aβ40] = [IAT]

[Aβ48] = [AICD 49-99] – [VIT]
[Aβ45] = [VIT] – [TVI]
[Aβ42 ]= [TVI] – [VVIA]
[Aβ38] = [VVIA]

**b**

| | Aβ49 | Aβ48 | Aβ46 | Aβ45 | Aβ43 | Aβ42 | Aβ40 | Aβ38 |
|---|---|---|---|---|---|---|---|---|
| WT | 90 | 118 | 45 | 60 | 10 | 132 | 193 | 62 |
| P117L | 151 | 172 | 20 | -7 | 20 | 28 | 24 | 16 |
| I143T | 9 | 32 | 22 | 0 | 0 | nd | nd | nd |
| L166P | 9 | 19 | 24 | 0 | 0 | nd | nd | nd |
| G384A | 46 | 126 | 20 | -6 | 19 | 25 | 24 | 17 |
| L435F | -5 | 29 | 28 | 0 | 0 | nd | nd | nd |
| L286V | -8 | 35 | 33 | 21 | 44 | 62 | 80 | 26 |

**c**

| | Total Aβs | Total AICD | Aβ40+Aβ43+Aβ46+Aβ49 | AICD 50-99 | Aβ38+Aβ42+Aβ45+Aβ48 | AICD 49-99 |
|---|---|---|---|---|---|---|
| **WT** | 714 | 668 | 340 | 334 | 374 | 334 |
| **P117L** | 427 | 434 | 217 | 204 | 210 | 230 |
| **I143T** | 64 | 51 | 32 | 23 | 32 | 28 |
| **L166P** | 52 | 50 | 33 | 25 | 19 | 26 |
| **G384A** | 274 | 289 | 111 | 115 | 163 | 174 |
| **L435F** | 52 | 47 | 23 | 24 | 29 | 23 |
| **L286V** | 297 | 301 | 151 | 141 | 146 | 160 |

**Extended Data Figure 4. Quantification of levels of each Aβ product from processing of APP substrate by WT vis-à-vis FAD-mutant γ-secretase. a,** Calculation of concentration of each Aβ species produced by incubation of purified γ-secretase complexes and C100Flag, based on concentrations of co-products. **b,** Concentration of each Aβ species in nM produced from reaction mixtures of 3 μM of C100Flag by 30 nM WT or FAD-mutant γ-secretase incubated at 37 °C for 16 h. Where the net level of precursor Aβ peptide for a given trimming step is zero, the net level of the trimmed Aβ product was not determined (nd). **c,** Total Aβ levels equal total AICD levels for each γ-secretase variant. Equimolar levels are also observed for Aβ peptides along the Aβ40 and Aβ42 pathways compared to corresponding AICD species (AICD50-99 and AICD49-99, respectively).

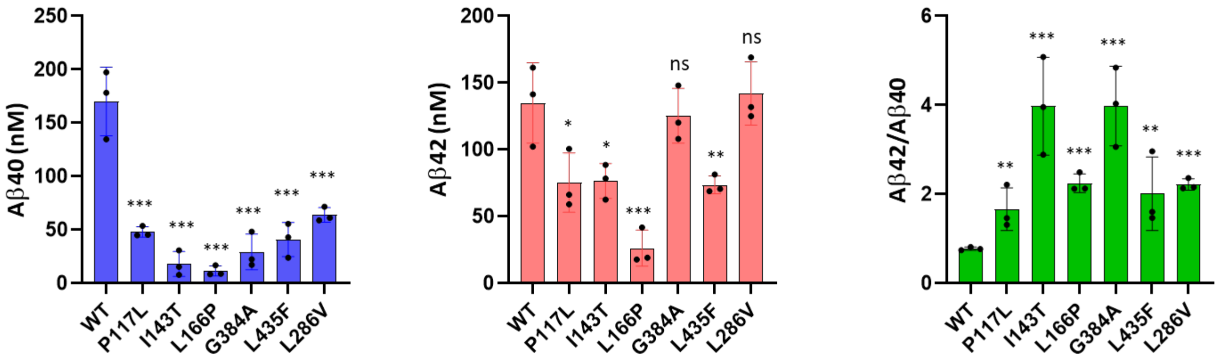

**Extended Data Figure 5. Aβ40, Aβ42, and Aβ42/Aβ40 ratio from cleavage of APP substrate by WT and FAD-mutant γ-secretases.** Aβ40 and Aβ42 produced by incubation of purified γ-secretase and C100Flag, as determined by specific ELISAs, and the resulting Aβ42/Aβ40 ratios. N=3, unpaired two-tailed T-test of FAD mutants compared to WT, *$p \leq 0.05$, **$p \leq 0.01$, ***$p \leq 0.001$.

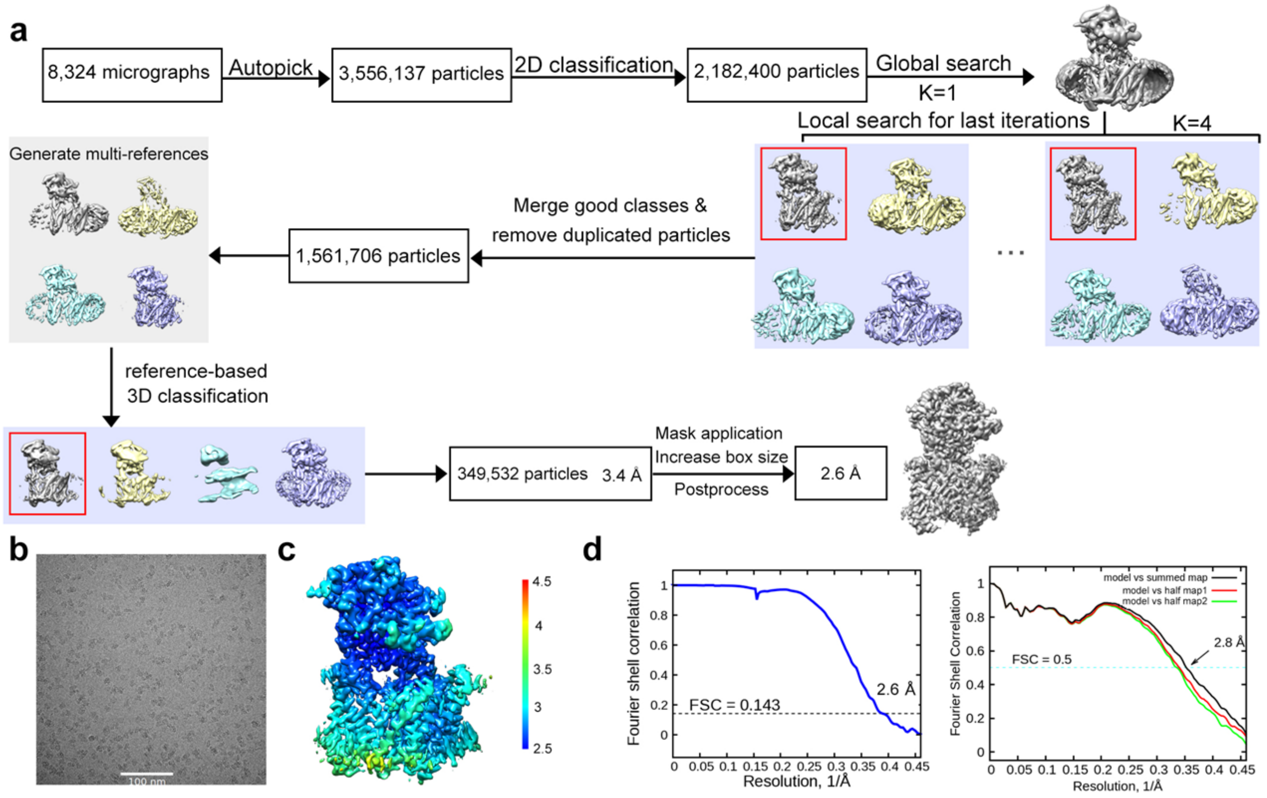

**Extended Data Figure 6. Cryo-EM analysis of SB-250 bound to human γ-secretase. a,** A flowchart of cryo-EM data processing. Please refer to Material and Methods for details. **b,** Representative cryo-EM micrograph of SB-250 bound human γ-secretase. A scale bar of 100 nm is shown. **c**, Color-coded local resolution distribution in Å of the final reconstruction, estimated by RELION-3.0. **d**, The final average resolution of the reconstruction of SB-250 bound human γ-secretase is 2.6 Å based on the 0.143 FSC curve. (left panel) The FSC curves of the refined model versus the maps that it is refined against (black); the model refined in the first of the two independent maps used for the FSC calculation versus that same map (red); and the model refined in the first of the two independent maps versus the second independent map (green) (right panel). The small difference between the red and green curves indicates that the refinement did not suffer from overfitting.

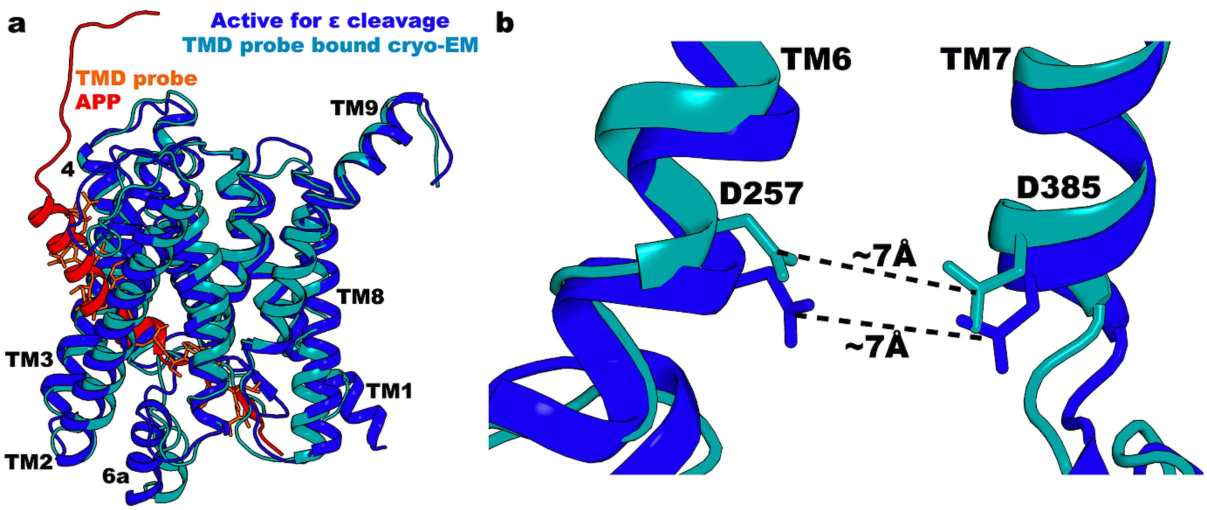

**Extended Data Fig. 7. Overlap of cryoEM structure of γ-secretase with TMD mimetic SB-250 and molecular dynamics simulation of activated γ-secretase poised for intramembrane proteolysis of APP substrate. a,** Overlap of PSEN1 and APP-C83/**SB-250**. **b,** Overlap of active site with catalytic aspartates D257 and D385.

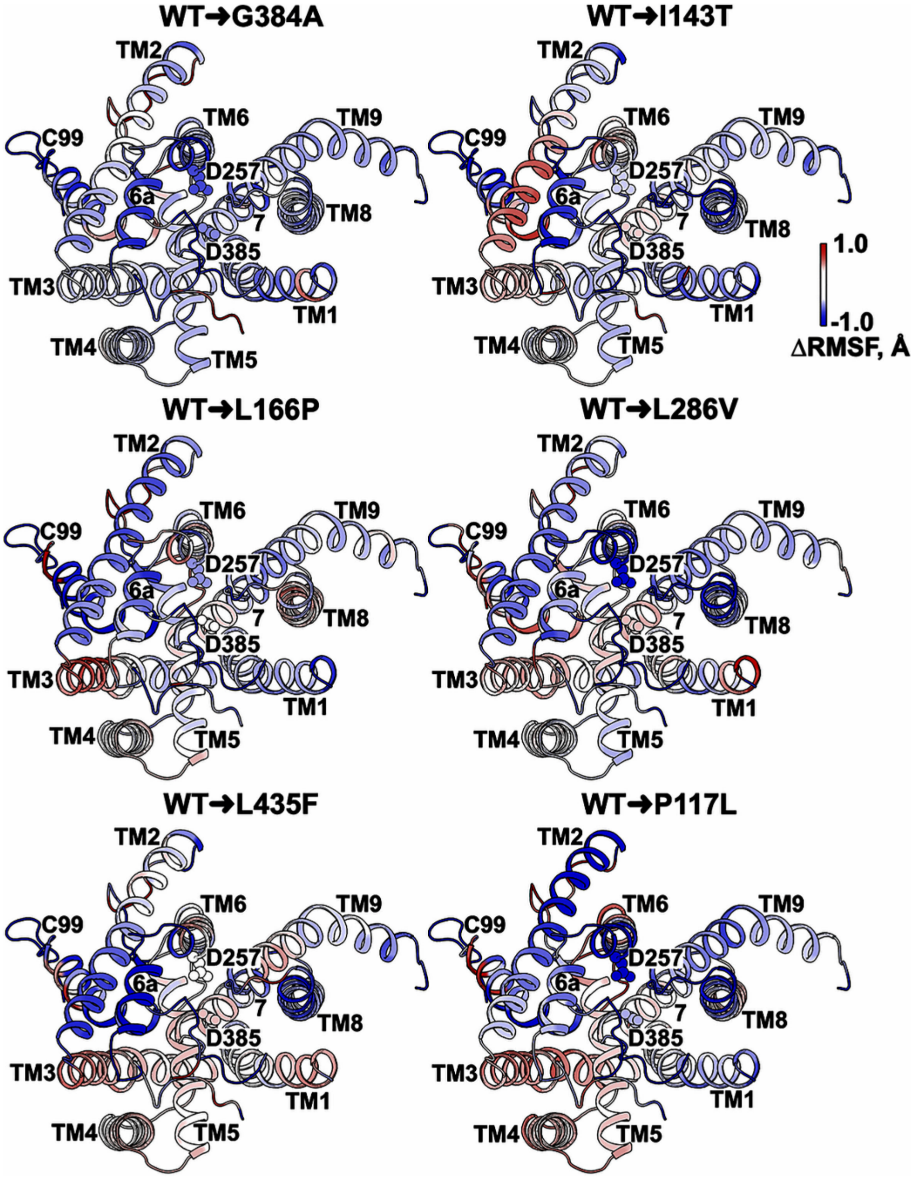

**Extended Data Fig. 8. GaMD simulations of γ-secretase bound to C99 substrate show overall reduced flexibility with FAD PSEN1 mutations, implying complex stabilization.** Images are as in main Fig. 2f but with a 90° rotation, looking at E-S complexes from the cytosolic side, to provide a better view of the active site.

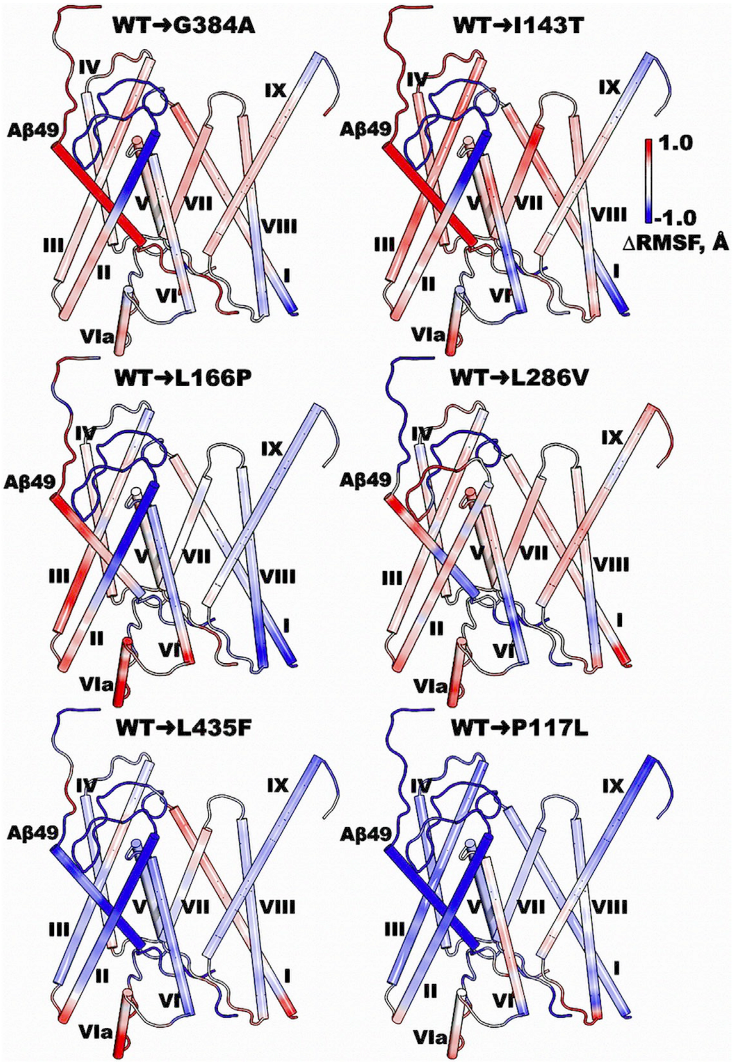

**Extended Data Fig. 9. Molecular dynamics simulations of γ-secretase bound to Aβ49**. Differences in RMSF between WT and FAD-mutant γ-secretase bound to Aβ49 show the FAD-mutant PSEN1 P117L leads to the most reduced conformational flexibility, especially for bound Aβ49, implying enzyme-intermediate complex stabilization.

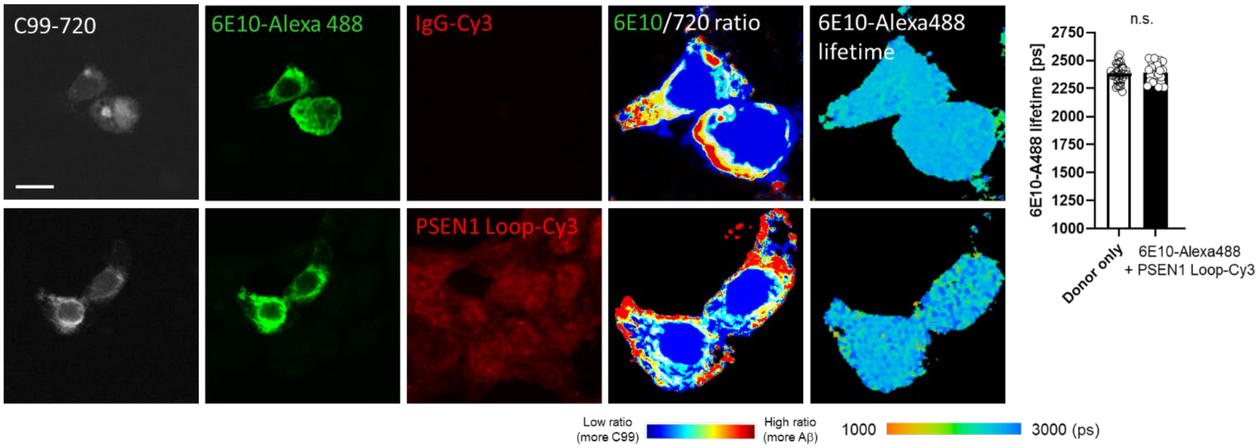

**Extended Data Fig. 10.** Control FLIM experiments using quencher-antibody that bind distal to 6E10 binding site on C99/Aβ. Specifically, rabbit monoclonal antibody EP2000Y to PSEN1 Loop was used in place of anti-nicastrin antibody NBP2-57365 used in main Fig. 3.

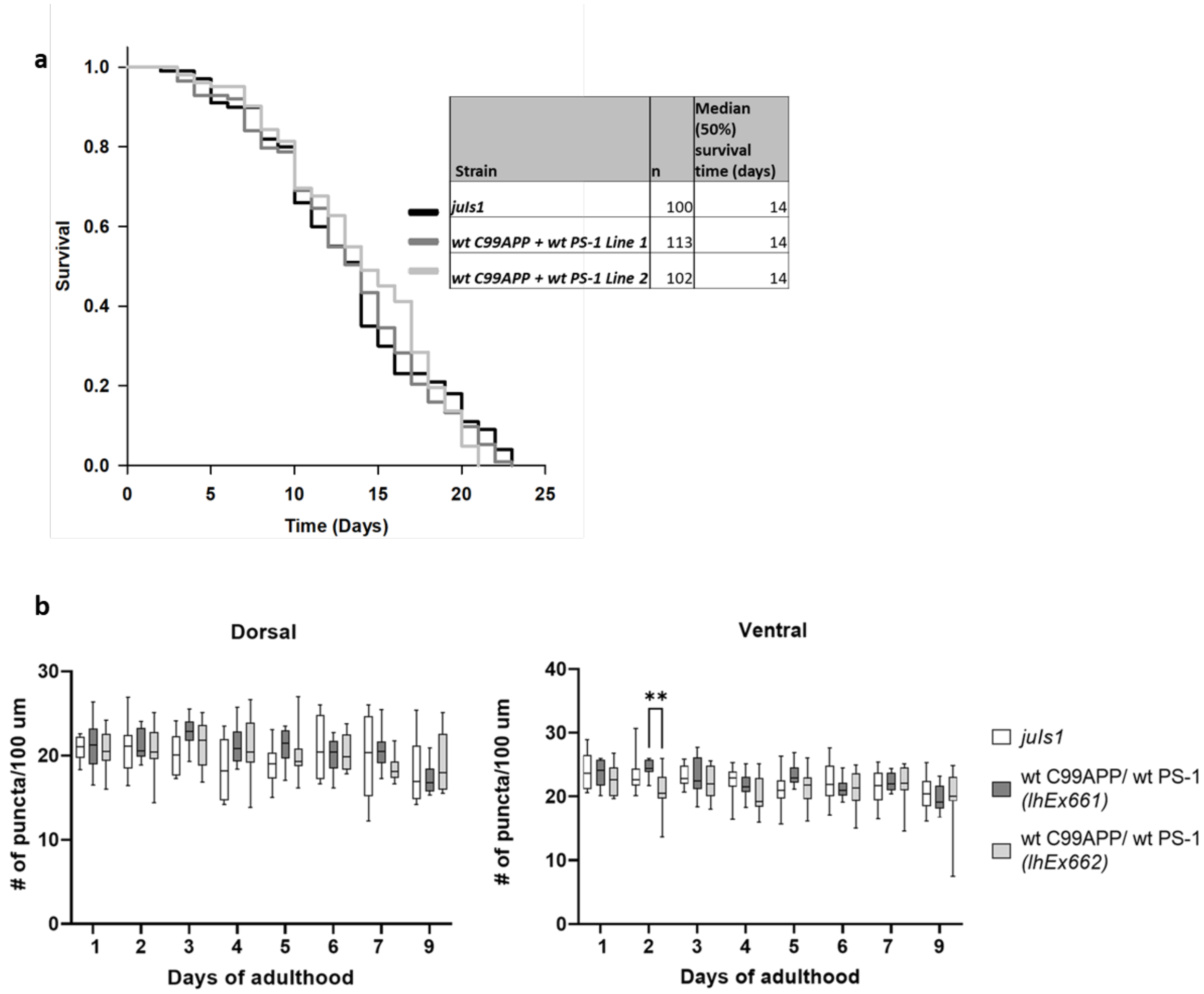

**Extended Data Fig. 11. Neuronal co-expression of WT C99 + WT PSEN1 does not affect *C. elegans* lifespan or synaptic puncta number. a**, Life span of parental line *juIs1* versus two double transgenic lines C99 + PSEN1. Kaplan-Meier curves of n animals show the fraction of animals alive at different days. **b**, Quantification of the dorsal and ventral synaptic puncta in the transgenic lines. Average number of synaptic puncta per 100 $\mu$m in n transgenic worms for each day is shown in the vertical box plots. Horizontal lines at the top and bottom of each box represent the maximum and minimum values, respectively. Upper and lower ends of a box mark quartiles Q1 and Q3 values, respectively. The horizontal line inside the box shows the median value and marks Q2). Two-way ANOVA for all possible pairs using Tukey's post hoc test, $**p \leq 0.01$. Note that the only statistically significant difference observed was between the two double transgenic lines on day 3 in the ventral nerve cord. No significant differences were observed between either double transgenic line and the parental line, on any day in either nerve cord.

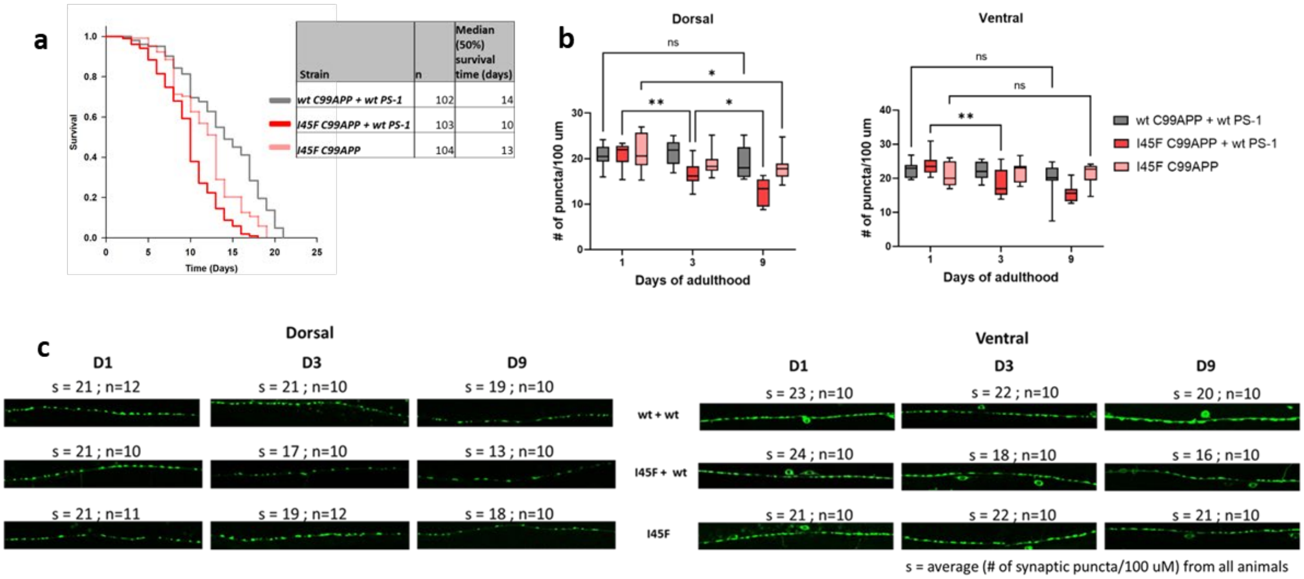

**Extended Data Fig. 12. Repeat of *C. elegans* experiments shown in main Fig. 4b-d with independent lines for all transgenes and transgene combinations. a**, Life span of double transgenic lines C99 + PSEN1 versus C99 I45F + PSEN1 and monogenic line C99 I45F. **b**, Quantification of the dorsal and ventral synaptic puncta in the transgenic lines. Two-way ANOVA for all possible pairs using Tukey's post hoc test, $*p \leq 0.05$, $**p \leq 0.01$, $***p \leq 0.001$, $****p < 0.0001$. Statistical differences shown only for days significance was seen within a given line. **c**, Representative 100 μm sections of confocal microscopic images dorsal (left) and ventral (right) nerve cords of transgenic animals. 's' denotes average number of synaptic puncta per 100 μm from n animals.

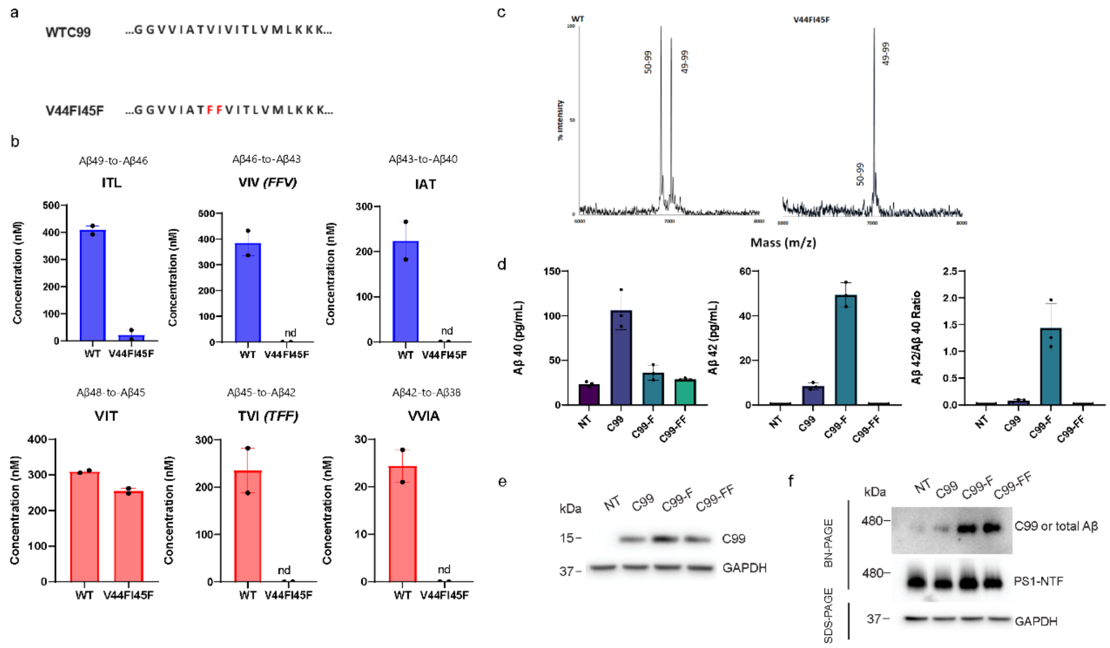

**Extended Data Fig. 13. V44F/I45F double mutation of APP substrate blocks Aβ46→Aβ43 and Aβ45→Aβ42 cleavage steps by γ-secretase and stabilizes E-S complexes. a,**Transmembrane domain sequence alignment of WT versus V44F/I45F C99. **b,** Concentration of tri- or tetra-peptides detected by LC-MS/MS analysis from WT C100-FLAG and V44F/I45F C100-FLAG. Cleavage resulting in Aβ40 production is represented by ITL→VIV→IAT (blue) and cleavage resulting in Aβ42 production is represented by VIT→TVI→VVIA (red). Cleavage steps from Aβ46→Aβ43 and Aβ45→Aβ42 for the V44F/I45F mutation produce tripeptides FFV and TFF, respectively. Notation "nd" indicates tri- or tetra-peptides unable to be detected by LC-MS/MS. 30 nM of enzyme and 5 μM of substrate were incubated at 37 °C for 16 h for both trials. The average and range of both trials is represented in the graph. **c,** MALDI-TOF mass spectrometric analysis of AICD 50-99 and AICD 49-99 produced by γ-secretase with WT versus V44F/I45F C100-FLAG. Note the V44F/I45F double mutation leads to substantially reduced relative AICD50-99 production. **d,** Secreted Aβ40 and Aβ42 levels in the culture media of HEK-293 cells stably expressing C99 and its mutants I45F and V44F/I45F were measured by specific ELISAs. Aβ42/Aβ40 ratios are also indicated. "NT" indicates the non-transfected (NT) control. Data are the mean ± S.E. (error bars) from three independent experiments. **e,** Cellular C99 levels in HEK-293 cells stably expressing C99 and its mutants were examined by Western blotting using monoclonal antibody 6E10. **f,** 10 μg of protein from the various HEK-293 cell lysates were subjected to BN-PAGE (1% digitonin) or SDS-PAGE and analyzed by Western blotting using antibodies specific for C99/Aβ, PS1-NTF and GAPDH. Data are representative of three independent experiments.

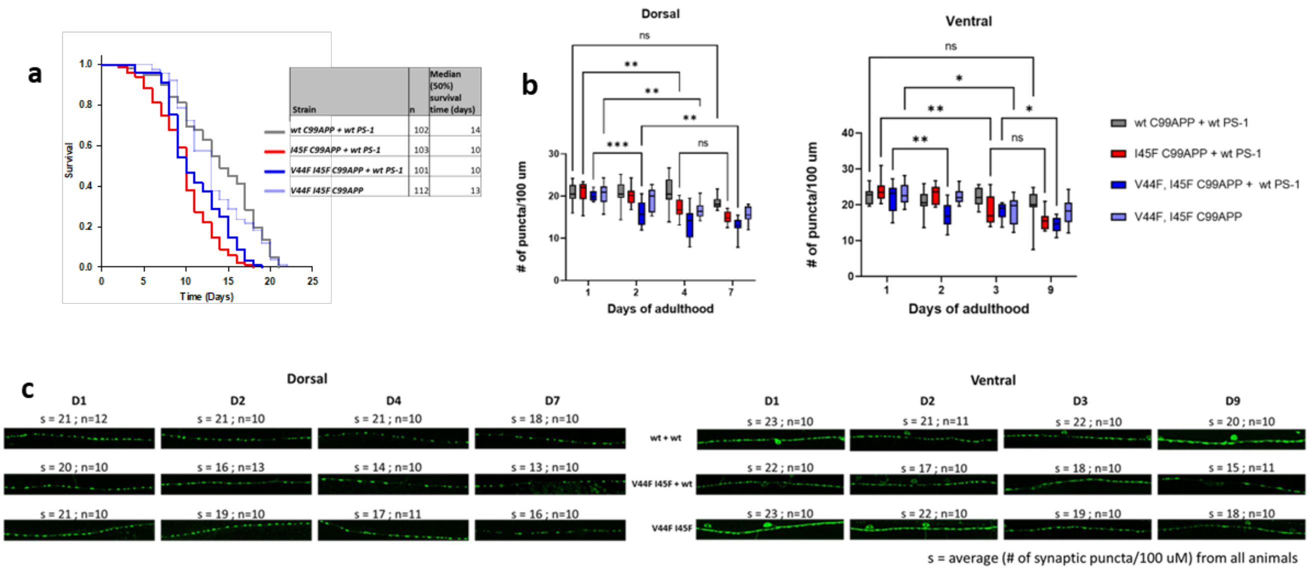

**Extended Data Fig. 14. Repeat of *C. elegans* experiments shown in main Fig. 4e-g with independent lines for all transgenes and transgene combinations. a**, Life span of double transgenic line C99 V44F/I45F + PSEN1 and monogenic line C99 V44F/I45F compared with C99 + PSEN1 and C99 I45F + PSEN1 lines. **b**, Quantification of dorsal and ventral synaptic puncta in these transgenic lines. Two-way ANOVA for all possible pairs using Tukey's post hoc test, *p≤ 0.05, **p≤ 0.01, ***p≤ 0.001, ****p< 0.0001. Statistical differences shown only for earliest day significance was seen from Day 1 in a given line. **c**, 100 μm sections of representative confocal microscopic images of dorsal and ventral synaptic puncta in these transgenic lines. The same data used in Extended Data Fig. 12 for C99 + PSEN1 and C99 I45F + PSEN1 were reproduced here for direct comparison with C99 V44F/I45F + PSEN1 and C99 V44F/I45F.

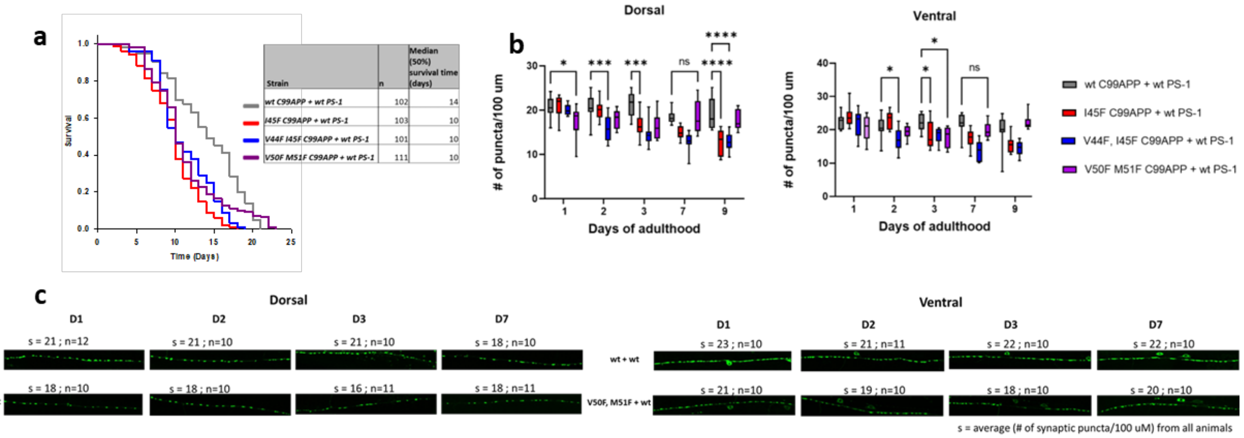

**Extended Data Fig. 15. Repeat of *C. elegans* experiments shown in main Fig. 5a-c with independent lines for all transgenes and transgene combinations. a**, Life span analysis of double transgenic line C99 V50F/M51F + PSEN1 compared with C99 + PSEN1, C99 I45F + PSEN1 and C99 V44F/I45F + PSEN1 lines. **b**, Quantification of dorsal and ventral synaptic puncta in these transgenic lines. Two-way ANOVA for all possible pairs using Tukey's post hoc test, *p≤ 0.05, **p≤ 0.01, ***p≤ 0.001, ****p< 0.0001. **c**, 100 μm sections of representative confocal microscopic images of dorsal and ventral synaptic puncta in these transgenic lines. The same data used in Extended Data Fig. 12 and 14 for C99 + PSEN1, C99 I45F + PSEN1 and C99 V44F/I45F + PSEN1 were reproduced here for direct comparison with C99 V50F/M51F + PSEN1.

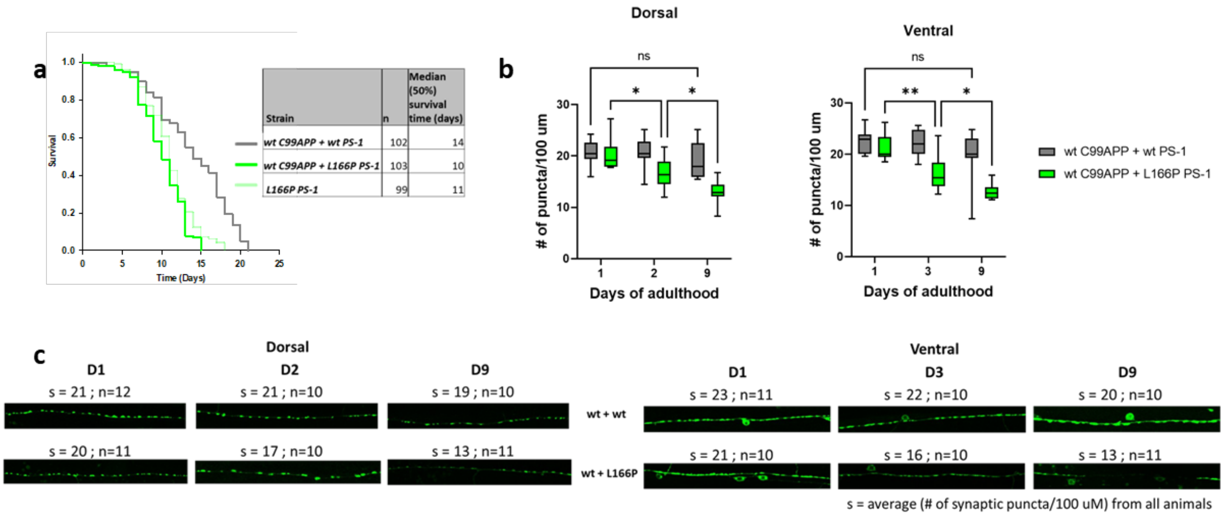

**Extended Data Fig. 16. Repeat of *C. elegans* experiments shown in main Fig. 5d-f with independent lines for all transgenes and transgene combinations. a**, Life span analysis of double transgenic line C99 + PSEN1 L166P and single transgenic line PSEN1 L166P compared with C99 + PSEN1. **b**, Quantification of dorsal and ventral synapses in double transgenic lines C99 + PSEN1 and C99 + PSEN1 L166P. Two-way ANOVA for all possible pairs using Tukey's post hoc test, $*p \leq 0.05$, $**p \leq 0.01$, $***p \leq 0.001$, $****p < 0.0001$. *Note*: L166P PS-1 monogenic lines were sensitive to the anesthetic agent, phenoxy propanol, leading to dying on agarose pads during sample preparation for imaging. Thus, confocal imaging of these animals was not possible. **c**,100 μm sections of representative fluorescence microscopic images of dorsal and ventral synaptic puncta in these transgenic lines. The same data used in Fig. 12 for C99 + PSEN1 were reproduced here for direct comparison with C99 + PSEN1 L166P and PSEN1 L166P (lifespan) or with only C99 + PSEN1 L166P (synaptic puncta).

**Extended Data Table 1**
**Cryo-EM data collection, refinement and validation statistics**

| | #1 name (EMDB-36948) (PDB 8K8E) |
|---|---|
| **Data collection and processing** | |
| Magnification | 81,000 |
| Voltage (kV) | 300 |
| Electron exposure (e–/Å$^2$) | 50 |
| Defocus range (μm) | ~1.5-~1.8 |
| Pixel size (Å) | 1.0825 |
| Symmetry imposed | C1 |
| Initial particle images (no.) | 3,556,137 |
| Final particle images (no.) | 349,532 |
| Map resolution (Å) | 2.6 |
| FSC threshold | 0.143 |
| Map resolution range (Å) | 2.5-4 |
| **Refinement** | |
| Initial model used (PDB code) | 6IYC |
| Model resolution (Å) | 2.8 |
| FSC threshold | 0.5 |
| Model resolution range (Å) | 2.8-256 |
| Map sharpening *B* factor (Å$^2$) | -10 |
| Model composition | |
| Non-hydrogen atoms | 10,856 |
| Protein residues | 1,324 |
| Ligands | 26 |
| *B* factors (Å$^2$) | |
| Protein | 101.20 |
| Ligand | 126.72 |
| R.m.s. deviations | |
| Bond lengths (Å) | 0.004 |
| Bond angles (°) | 0.621 |
| Validation | |
| MolProbity score | 1.98 |
| Clashscore | 7.36 |
| Poor rotamers (%) | 1.77 |
| Ramachandran plot | |
| Favored (%) | 94.30 |
| Allowed (%) | 5.70 |
| Disallowed (%) | 0.00 |

#### Extended Data Table 2
#### Survival curve analysis of parental *C. elegans* line *juIs1* versus transgenic lines expressing WT C99 + WT PSEN1 in neurons.

| Transgenic lines | Total # of worms (n) | Survival Time (Days) | Std. Error | Survival Time (Days) | Std. Error | Survival Time (Days) | Std. Error | Survival Time (Days) | Std. Error |
|---|---|---|---|---|---|---|---|---|---|
| | | **Mean** | | **25th Percentile[a]** | | **50th Percentile[b]** | | **75th Percentile[c]** | |
| ***Extended data fig 11*** | | | | | | | | | |
| *juIs1* | 100 | 13.34 | 0.515 | 16 | 1.403 | 14 | 0.572 | 10 | 0.592 |
| *wt C99APP + wt PS-1 1st line (lhEx661)* | 113 | 13.274 | 0.476 | 17 | 0.611 | 14 | 0.659 | 10 | 0.867 |
| *wt C99APP + wt PS-1 2nd line (lhEx662)* | 102 | 13.971 | 0.472 | 18 | 0.365 | 14 | 0.918 | 10 | 0.387 |

Statistical analysis of Kaplan-Meier survival curves showed no significant difference between any of the curves (P = 0.851). [a]survival time where 75% of the animals are expected to survive. [b]median survival time. [c]survival time where 25% of the animals are expected to survive.

**Extended Data Table 3**

**Survival curve analysis of *C. elegans* transgenic lines using data from experiments shown in main Figs 4 and 5.**

| **Transgenic lines** | **Total # of worms (n)** | **Survival Time (Days)** | **Std. Error** | **Survival Time (Days)** | **Std. Error** | **Survival Time (Days)** | **Std. Error** | **Survival Time (Days)** | **Std. Error** |
|---|---|---|---|---|---|---|---|---|---|
| | | **Mean** | | **25th Percentile[a]** | | **50th Percentile[b]** | | **75th Percentile[c]** | |
| ***Figure 4b*** | | | | | | | | | |
| *wt C99APP + wt PS-1* | 113 | 13.274 | 0.476 | 17 | 0.611 | 14 | 0.659 | 10 | 0.867 |
| *I45F C99APP + wt PS-1* | 99 | 10.02 | 0.368 | 13 | 0.548 | 10 | 0.48 | 7 | 0.269 |
| *I45F C99APP* | 100 | 12.97 | 0.439 | 16 | 0.481 | 14 | 0.762 | 10 | 0.612 |
| ***Figure 4e*** | | | | | | | | | |
| *wt C99APP + wt PS-1* | 113 | 13.274 | 0.476 | 17 | 0.611 | 14 | 0.659 | 10 | 0.867 |
| *I45F C99APP + wt PS-1* | 99 | 10.02 | 0.368 | 13 | 0.548 | 10 | 0.48 | 7 | 0.269 |
| *V44F I45F C99APP + wt PS-1* | 102 | 9.51 | 0.335 | 11 | 0.408 | 9 | 0.404 | 7 | 0.422 |
| *V44F I45F C99APP* | 102 | 11.961 | 0.293 | 14 | 0.367 | 12 | 0.488 | 10 | 0.302 |
| ***Figure 5a*** | | | | | | | | | |
| *wt C99APP + wt PS-1* | 113 | 13.274 | 0.476 | 17 | 0.611 | 14 | 0.659 | 10 | 0.867 |
| *I45F C99APP + wt PS-1* | 99 | 10.02 | 0.368 | 13 | 0.548 | 10 | 0.48 | 7 | 0.269 |
| *V44F I45F C99APP + wt PS-1* | 102 | 9.51 | 0.335 | 11 | 0.408 | 9 | 0.404 | 7 | 0.422 |
| *V50F M51F C99APP + wt PS-1* | 99 | 11.343 | 0.377 | 14 | 0.814 | 11 | 0.613 | 9 | 0.378 |
| ***Figure 5d*** | | | | | | | | | |
| *wt C99APP + wt PS-1* | 113 | 13.274 | 0.476 | 17 | 0.611 | 14 | 0.659 | 10 | 0.867 |
| *wt C99APP + L166P PS-1* | 110 | 10.336 | 0.346 | 13 | 0.14 | 11 | 0.583 | 8 | 0.752 |
| *L166P PS-1* | 99 | 10.343 | 0.348 | 13 | 0.64 | 9 | 0.553 | 8 | 0.159 |

Statistical analysis of Kaplan-Meier survival curves. [a]survival time where 75% of the animals are expected to survive. [b]median survival time. [c]survival time where 25% of the animals are expected to survive.

**Extended Data Table 4**

**Pairwise comparison of survival curves of *C. elegans* transgenic lines using data from experiments shown in main Figs 4 and 5.**

| Pairwise multiple comparison (Holm-Sidak method) | Overall Significance level= 0.05 | |
|---|---|---|
| **Comparisons** | **P Value** | **Significant?** |
| ***Figure 4b*** | | |
| *wt C99APP + wt PS-1* **vs** *I45F C99APP + wt PS-1* | 3.46E-09 | Yes |
| *I45F C99APP + wt PS-1* **vs** *I45F C99APP* | 1.21E-08 | Yes |
| *wt C99APP + wt PS-1* **vs** *I45F C99APP* | 0.151 | No |
| ***Figure 4e*** | | |
| *wt C99APP + wt PS-1* **vs** *V44F I45F C99APP + wt PS-1* | 1.43E-11 | Yes |
| *wt C99APP + wt PS-1* **vs** *V44F I45F C99APP* | 0.000187 | Yes |
| *V44F I45F C99APP + wt PS-1* **vs** *V44F I45F C99APP* | 2.31E-05 | Yes |
| *I45F C99APP + wt PS-1* **vs** *V44F I45F C99APP + wt PS-1* | 0.211 | No |
| ***Figure 5a*** | | |
| *wt C99APP + wt PS-1* **vs** *V50F M51F C99APP + wt PS-1* | 0.00066 | Yes |
| *I45F C99APP + wt PS-1* **vs** *V50F M51F C99APP + wt PS-1* | 0.0217 | Yes |
| *V44F I45F C99APP + wt PS-1* **vs** *V50F M51F C99APP +wt PS-1* | 0.00092 | Yes |
| ***Figure 5d*** | | |
| *wt C99APP + wt PS-*1 **vs** *wt C99APP + L166P PS-1* | 3.7E-09 | Yes |
| *wt C99APP + wt PS-1* **vs** *L166P PS-1* | 1.8E-07 | Yes |
| *L166P PS-1* **vs** *wt C99APP + L166P PS-1* | 0.823 | No |

**Extended Data Table 5**

**Survival curve analysis of *C. elegans* transgenic lines using data from experiments shown in Extended Data Fig. 12, 14-16.**

| Transgenic lines | Total # of worms (n) | Survival Time (Days) | Std. Error | Survival Time (Days) | Std. Error | Survival Time (Days) | Std. Error | Survival Time (Days) | Std. Error |
|---|---|---|---|---|---|---|---|---|---|
| | | Mean | | 25th Percentile[a] | | 50th Percentile[b] | | 75th Percentile[c] | |
| ***Extended data fig. 12a*** | | | | | | | | | |
| *wt C99APP + wt PS-1* | 102 | 13.971 | 0.472 | 18 | 0.365 | 14 | 0.918 | 10 | 0.387 |
| *I45F C99APP + wt PS-1* | 103 | 9.806 | 0.334 | 12 | 0.528 | 10 | 0.224 | 7 | 0.63 |
| *I45F C99APP* | 104 | 11.827 | 0.38 | 14 | 0.409 | 13 | 0.319 | 8 | 0.513 |
| ***Extended data fig. 14a*** | | | | | | | | | |
| *wt C99APP + wt PS-1* | 102 | 13.971 | 0.472 | 18 | 0.365 | 14 | 0.918 | 10 | 0.387 |
| *I45F C99APP + wt PS-1* | 103 | 9.806 | 0.334 | 12 | 0.528 | 10 | 0.224 | 7 | 0.63 |
| *V44F I45F C99APP + wt PS-1* | 101 | 11.158 | 0.355 | 14 | 0.566 | 10 | 0.442 | 9 | 0.271 |
| *V44F I45F C99APP* | 112 | 13.277 | 0.387 | 16 | 1.051 | 13 | 0.466 | 10 | 0.364 |
| ***Extended data fig. 15a*** | | | | | | | | | |
| *wt C99APP + wt PS-1* | 102 | 13.971 | 0.472 | 18 | 0.365 | 14 | 0.918 | 10 | 0.387 |
| *I45F C99APP + wt PS-1* | 103 | 9.806 | 0.334 | 12 | 0.528 | 10 | 0.224 | 7 | 0.63 |
| *V44F I45F C99APP + wt PS-1* | 101 | 11.158 | 0.355 | 14 | 0.566 | 10 | 0.442 | 9 | 0.271 |
| *V50F M51F C99APP + wt PS-1* | 111 | 11.252 | 0.422 | 13 | 0.695 | 10 | 0.318 | 8 | 0.597 |
| ***Extended data fig. 16a*** | | | | | | | | | |
| *wt C99APP + wt PS-1* | 102 | 13.971 | 0.472 | 18 | 0.365 | 14 | 0.918 | 10 | 0.387 |
| *wt C99APP + L166P PS-1* | 103 | 10.107 | 0.288 | 13 | 0.194 | 10 | 0.423 | 8 | 0.391 |
| *L166P PS-1* | 99 | 11.04 | 0.309 | 13 | 0.399 | 11 | 0.273 | 9 | 0.517 |

Statistical analysis of Kaplan-Meier survival curves. [a]survival time where 75% of the animals are expected to survive. [b]median survival time. [c]survival time where 25% of the animals are expected to survive.

**Extended Data Table 6**

**Pairwise comparison of survival curves of *C. elegans* transgenic lines using data from experiments shown in Extended Data Fig. 12, 14-16.**

| Pairwise multiple comparison (Holm-Sidak method) | Overall significance level = 0.05 | |
|---|---|---|
| **Comparisons** | **P Value** | **Significant?** |
| ***Extended data fig. 12a*** | | |
| *wt C99APP + wt PS-1 vs. I45F C99APP + wt PS-1* | 1.11E-13 | Yes |
| *I45F C99APP + wt PS-1 vs I45F C99APP* | 1.61E-05 | Yes |
| *wt C99APP + wt PS-1 vs. I45F C99APP* | 2.72E-05 | Yes |
| ***Extended data fig. 14a*** | | |
| *wt C99APP + wt PS-1 vs. V44F I45F C99APP + wt PS-1* | 4.53E-08 | Yes |
| *wt C99APP + wt PS-1 vs. V44F I45F C99APP* | 0.239 | No |
| *V44F I45F C99APP + wt PS-1 vs V44F I45F C99APP* | 4.91E-05 | Yes |
| *I45F C99APP + wt PS-1 vs V44F I45F C99APP + wt PS-1* | 0.0156 | Yes |
| ***Extended data fig. 15a*** | | |
| *wt C99APP + wt PS-1 vs. V50F M51F C99APP + wt PS-1* | 0.00897 | Yes |
| *I45F C99APP + wt PS-1 vs V50F M51F C99APP + wt PS-1* | 0.0176 | Yes |
| *V44F I45F C99APP + wt PS-1 vs V50F M51F C99APP +wt PS-1* | 0.481 | No |
| ***Extended data fig. 16a*** | | |
| *wt C99APP + wt PS-1 vs. wt C99APP + L166P PS-1* | 7.26E-14 | Yes |
| *wt C99APP + wt PS-1 vs. L166P PS-1* | 2.3E-09 | Yes |
| *L166P PS-1 vs. wt C99APP + L166P PS-1* | 0.0381 | Yes |
